# Plasticity in ear density drives complementarity effects and yield benefits in wheat variety mixtures

**DOI:** 10.1101/2025.02.25.640070

**Authors:** Laura Stefan, Nathalie Colbach, Dario Fossati, Silvan Strebel, Lilia Levy Häner

**Affiliations:** Cultural Techniques and Varieties in Arable Farming, Agroscope, Nyon, Switzerland; Agroécologie, INRAE, Institut Agro, Université Bourgogne Europe, 21000 Dijon, France; Field-Crop Breeding and Genetic Resources, Agroscope, Nyon, Switzerland

## Abstract

Variety mixtures represent a promising option to sustainably increase the productivity of grain cropping systems, but the underlying processes driving potential yield benefits remain poorly understood. Notably, the role of variety-specific phenotypic changes in mixtures – defined here as plasticity – and their effects on plant-plant interactions has scarcely been evaluated. Here, we examined the trait responses of 8 Swiss wheat varieties when grown in mixtures, and how these plastic changes contributed to overyielding, complementarity and selection effects. For this, we conducted an outdoor field experiment in 3 years and 3 sites, where wheat varieties were grown in 2-way mixtures and in pure stands. We used a visual criterion (awns) to differentiate individuals of the different varieties in mixtures. We found significant plastic changes in response to mixing for several traits in 7 varieties. Furthermore, mixture-induced plasticity in ear density was the main driver of overyielding, itself largely dominated by complementarity effects. An additional experiment allowed us to positively link plasticity in ear density to the speed of tillering onset under shading. This study improves our understanding of the plastic processes fostering overyielding in variety mixtures, and provides a key criterion – tillering onset under shade – as a potential breeding target of cultivars for mixtures.

## Introduction

The main agronomical challenge of the current and next decades is to make food production more sustainable and less detrimental to the environment, while maintaining – or even increasing – crop yield in order to feed an ever-growing population (Tilman *et al*., 2011; Weiner, 2017). With this objective in mind, bringing biodiversity back to the agricultural fields is more relevant than ever (Montoya *et al*., 2020; Brooker *et al*., 2023). At the field scale, biodiversity indeed promotes and stabilizes ecosystem services, such as yield, nutrient cycling, pollination, and pests and diseases limitation (Tamburini *et al*., 2020; Cappelli *et al*., 2022). In this context, variety mixtures represent a practical way to integrate diversity into agricultural fields, as they do not require technical or mechanical adjustments (Wuest *et al*., 2021). In the case of wheat cultivation, variety mixtures have indeed shown significant potential to sustainably maintain or even increase grain biomass per area while improving the stability of grain crop production (Borg *et al*., 2017; Stefan *et al*., 2024). However, the mechanisms underlying these effects remain evasive (Li *et al*., 2023); this makes it difficult to understand which varieties to combine for yield benefits, and further prevents the adoption of variety mixtures.

In pure stands, decades of research and experiments in wheat cultivation have allowed to identify key traits for breeding high-performing varieties (Donald, 1968). Selection has notably targeted performance at the group (i.e. plot) level rather than at the individual (i.e. plant) level, thereby favouring varieties with low competitive abilities that could be grown at very high densities (Denison, 2012; Weiner, 2017). In mixtures however, the ecological processes governing plant-plant interactions between different varieties are not the same as in monocultures (Su *et al*., 2023), with the consequences that plants grow differently when surrounded by kin vs. non-kin neighbours (Dahlin *et al*., 2020). Therefore, the ideotypes and key traits identified for high performance in monocultures may not be optimal for mixtures (Barot *et al*., 2017; Kopp *et al*., 2023). Assessing how key plant traits change in mixtures compared to monocultures – i.e. plasticity in response to mixtures – is thus crucial to better understand plant-plant interactions in mixtures and to identify ideotypes for cultivation in mixtures.

In diverse plant communities, plant-plant interactions between species and/or varieties have been shown to shift to increased niche differentiation and/or facilitation (Wright *et al*., 2021; Stefan *et al*., 2022). In the framework of biodiversity-ecosystem functioning relationships, these two processes are encompassed in the Complementarity Effects (Loreau & Hector, 2001). Complementarity notably occurs when varieties use the resources differently, compete less intensely, and therefore are more productive than their corresponding pure stands (Chesson, 1994; Zuppinger-Dingley *et al*., 2014). To illustrate this process with an example, in intercropping systems complementarity can typically occur when combining a cereal with a legume species, as the legume can use another source of nitrogen and therefore will not deprive the cereal from access to this resource in the soil (Jensen *et al*., 2020). In addition, diverse plant communities can be more productive because of the Selection Effects, which reflect the increased chance of including a high-yielding variety that will also dominate performance in more variety-rich communities (Tilman *et al*., 1997; Loreau & Hector, 2001). In variety mixtures, the importance, frequency and mechanisms of complementarity vs. selection effects are not fully elucidated, mostly because often the varieties’ individuals/grains cannot easily be distinguished from each other, which makes it almost impossible to determine the yield of individual cultivars in a mixture (but see (Gawinowski *et al*., 2024) with an extensive experimental design).

In this study, we used a visual trait (awns) to be able to distinguish and separate the individual varieties in 2-way mixtures. This allowed us to assess variety-specific trait changes from pure stands to mixtures and to calculate, for the first time to our knowledge, complementarity and selection effects in wheat variety mixtures.

The objectives were thus 3-fold: 1) to evaluate the plasticity from pure stands to mixtures of various wheat traits linked to productivity and grain quality for 8 Swiss wheat varieties contrasting for quality and yield potential; 2) to assess overyielding, complementarity, and selection effects in mixtures; 3) to understand the links between trait plasticity and overyielding, in order to find key traits relevant for selecting and breeding varieties for mixtures. This was done using field trials including 2-way mixtures – in which we could visually separate and harvest the varieties – and pure stands of the varieties, which took place in 3 sites during 3 growing seasons in Switzerland. In addition, because competition for light is the main resource for which plants compete in temperate arable cropping systems with inputs (Colbach *et al*., 2023; Gawinowski *et al*., 2024), we grew individuals of each variety as single plants in unshaded and shaded conditions to assess the phenological responses to shading. By improving our understanding of the processes underlying overyielding in mixtures, we wished to identify key traits that could be further used by breeders to select varieties for optimal cultivation in mixtures.

## Material and Methods

The study sites and experimental design were similar to the ones described in Stefan *et al*. (Stefan *et al*., 2025).

### Study sites

The study took place over the course of three growing seasons – 2020/2021, 2021/2022, and 2022/2023 – in three field sites across the Swiss Central Plateau. The three experimental sites were located in Changins (46°19′ N 6°14′ E, 455m a.s.l), Delley (46°55′ N 6°58′ E, 494m a.s.l) and Utzenstorf (47°97′ N 7°33′ E, 483m a.s.l.). The main differences between the field sites across the growing seasons were precipitations (794 mm in Delley, 756 mm in Utzenstorf, and 715 mm in Changins), and temperatures, with Utzenstorf being the coldest site (mean temperature of 8.7°C and minimum temperature of -2.1°C), followed by Delley (9°C and 0.2°C for mean and minimum temperatures). Changins was the warmest site, with a mean temperature of 10.3°C and a minimum temperature of 1.1°C. Detailed weather records can be found in Figures S1-S3.

### Field trials

We used 8 Swiss varieties as described in Stefan *et al*. (2024). The varieties included CH111.16373, CH211.14074, Bodeli, Campanile, Colmetta, Falotta, Molinera and Schilthorn. The varieties came from the Swiss national breeding program in 2020, and were chosen to include differences in yield potential, protein content, foliage shape, and awness, with no more than 15 cm difference in height and no more than 5 days difference in phenology at maturity. Five of them were awned (CH211.14074, Bodeli, Colmetta, Falotta, Molinera) and three of them lacked awns (CH111.16373, Campanile, Schilthorn). The experimental design consisted of all the varieties grown in pure stands, and all the possible 2-variety mixtures of one awned variety with an awnless variety, leading to a total of 15 mixtures. We focused on the 2-variety mixtures of one awned and one awnless variety in order to be able to visually distinguish and separate the varieties. Each pure stand and mixture were grown in plots of 7.05 m^2^ (1.5m × 4.7m). We replicated the experiment three times per site with the exact same variety and mixture composition, a complete randomized block design. Sowing density was 350 viable seeds/m^2^ for each pure stand and mixture plots. For the mixtures, seeds were mixed beforehand at a 2×50% mass ratio for 2-variety mixtures, e.g. 100g of variety A was mixed with 100g of variety B. This means that the sowing densities of each variety in mixture varied depending on the mixture, with the total sowing density remaining 350 seeds/m^2^. We chose this method of mixing as this is what is commonly done by farmers in Switzerland. Plots were sowed mechanically each autumn (see Table S1) and fertilized with ammonium nitrate at a rate of 140 N/ha in 3 applications on all sites (40 N/ha at tillering stage/BBCH 22–29; 60 N/ha at the beginning of stem elongation/BBCH 30–31; 40 N/ha at the flag leaf stage/BBCH 45–47). The trials were grown according to the Swiss *Extenso* scheme, i.e. without any fungicide, pesticide, or growth regulator. Weeds were regulated twice per season with the application of herbicides commonly used in Switzerland. Disease incidence was monitored throughout the growing seasons, but there were no major outbreaks in any of our field sites, and therefore disease incidence will not be presented here.

### Data acquisition

#### Plant height

In Changins, we measured plant height of 5 random individuals from each variety in each plot (i.e. in mixture plots and pure stands) at the phenology stage BBCH 69. Plant height was measured in centimeters, from the ground to the top of the ears, including awns.

#### Yield

At maturity, we manually harvested a strip of 30 cm × 1.5 m = 0.45 m^2^ along the width of each plot by cutting all individuals (i.e. stems with ears) within the strip right above the ground (see Fig. S4 for a schematic illustration). The harvested strip was located at more than 1 meter from the plot borders to avoid edge effect, and non-homogeneous parts of the plots were avoided. We subsequently air-dried the samples for two weeks. For pure stands samples, we counted the stems, weighted the total biomass, then threshed and weighted the grains. For mixture samples, we manually sorted the awned stems and ears from the awnless stems and ears. We counted the number of awned and awnless stems and ears in each sample and weighed the total biomass of the two varieties. We then separately threshed the awned and awnless individuals and subsequently weighed the grains, in order to obtain the yields from the awned variety and from the awnless variety.

#### Post-harvest analyses

Protein content (% of dry matter) was measured for each previously described samples (e.g. for each variety in pure stand, but also for each associated variety in mixture plots) with a near-infrared instrument (ProxiMateTM, Büchi instruments), and thousand kernel weight (TKW, g) with a Marvin seed analyzer (GTA Sensorik, Neubrandenburg, Germany).

### Light competition trial

In Changins, in 2021/2022, we grew single plants of each variety in full light and in shaded conditions to assess the phenological response of each variety to competition for light. The experiment took place in an outdoor agricultural field, where we covered the soil with a thin, dark, microperforated agricultural foil to avoid weed infestation. The agricultural foil contained one hole every 0.25m2, where varieties were grown as individual plants. Over half the soil surface, we constructed dome-shaped, metallic tunnels, which we covered with a shading net intercepting 66% of photosynthetically active radiation (PAR). Each variety was replicated 5 times per light treatment, with randomized blocks within each light treatment. The phenological development of each plant following the BBCH scale was regularly assessed (i.e. every 2-3 days).

### Data analyses

All the analyses were performed using R version 4.3.0 (R Core Team, 2019).

#### Plasticity

In order to assess how plants respond to being in mixtures, and whether this response is variety- or mixture-specific, we analysed the effects of variety and associated variety on trait changes in mixtures compared to pure stands. To do so, for the traits considered in this study – i.e. biomass (kg/0.45m^2^), grain yield (kg/0.45m^2^), TKW (g), protein content (%), harvest index (seed biomass (kg)/total above-ground biomass (kg)), ear density (number of ears – fertile tillers – per 0.45m^2^), and plant height (cm, in Changins only) – we calculated plasticity in mixtures of each variety by computing the difference between the measured trait value of the variety in the mixture and the measured trait value of the variety grown in pure stand. For traits whose calculation is independent of plant density – i.e. height, TKW, protein content and harvest index – this was calculated per variety and variety couple using the following formula:

> mixture-induced height plasticity of variety A in mixture (A,B) = height of variety A in mixture (A,B) – height of variety A in pure stand

For traits whose calculation was dependent on plant density – such as yield, biomass, and ear density – we calculated plasticity as the difference between the trait value of the variety in the mixture and the expected trait value based on the pure-stand value and the relative sowing density in the mixture, e.g.

> mixture-induced yield plasticity of variety A in mixture (A,B) = yield of variety A in mixture (A,B) – yield of variety A in pure stand*(sowing density of variety A in the mixture (A,B)/sowing density of variety A in pure stand)

Plasticity was computed per block, amounting to 3 replicate values per variety × associated variety × year × site.

#### General response of varieties

To investigate the response of trait plasticity at the variety level, we used linear mixed models with the function *lmer* from the R package *lme4* (Bates *et al*., 2015) with variety as fixed factor, and year, site, and replicate as well as variety combination as random factors. We included variety combination, year and site as random factors because we wanted to assess the response of varieties across environments and mixtures rather than environment-specific and mixture-specific responses.

#### Ordination analysis

The trait plasticity responses were assessed visually with a canonical analysis of principal coordinates (CAP; (Anderson & Willis, 2003)). For this, we took the average value over the three replicates per variety × associated variety × site × year for plasticity in biomass, yield, ear density, protein content, TKW, and harvest index. From this trait plasticity dataset, we calculated the associated distance matrix with Euclidean distance. The canonical analysis of principal coordinates was conducted on the trait plasticity distance matrix with the function *CAPdiscrim* from the *BiodiversityR* package (Kindt & Coe, 2005), with variety as an explanatory variable.

#### Mixture-specific response of varieties

To investigate the response of plasticity at the mixture-specific level (i.e., the response of each variety as a function of the associated variety), we created subsets of the dataset for each of the variety (e.g., a subset of all the plots containing Variety A). For each of these subsets, we fitted a linear mixed-effects model using *lmer* with variety combination as fixed factors, and year, site and replicate as random factors. This allowed us to assess the response of Variety A according to the associated variety, e.g. whether it is associated with variety B or variety C.

#### Overperformance, complementarity and selection effects

Here we aimed to assess whether mixtures were more or less productive than the respective pure stands, and how variety or variety combination played a role in these yield benefits. For yield and protein content, we thus calculated the corresponding overyielding and overperformance in protein content in mixtures as the difference between the observed and expected values of the mixtures, where the expected value is the sum of the values in pure stands weighted by the relative sowing density in the mixture (Loreau & Hector, 2001):

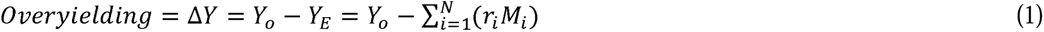

where N is the number of varieties in the plot, *r_i_* indicates the relative sowing density of the variety *i* in the mixture, and *M_i_* is the yield of the variety *i* in pure stand. Following the method developed by Loreau and Hector (2001), we partitioned overyielding into its two components, namely complementarity and selection effects:

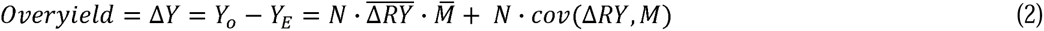

Here, N is the number of varieties in the plot; ΔRY (kg/kg) is the deviation from the expected relative yield of the variety in mixture, which is calculated as the ratio of observed relative yield of the variety in mixture to the yield of the variety in pure stand; 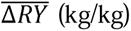 is the average of ΔRY of the varieties in the mixture. M (kg) is the yield of the variety in pure stand, and 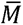 is the average of monoculture yields for the varieties in the mixture. The first component of the biodiversity effect equation 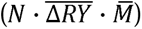 is the complementarity effect (CE, kg) and represents how much more individual varieties contribute to productivity than predicted from pure stands and the relative sowing densities in the mixture. The second component (N · *cov*(Δ*RY*, *M*)) is the selection effect (SE, kg) and describes the greater probability of more diverse communities to include highly productive varieties which also account for most of the productivity. These calculations were performed per variety combination × site × year × block.

The responses of overyielding, CE and SE to variety and variety combination were investigated using linear mixed-effects models, with presence/absence of each variety or variety combination as fixed factors and year, site, and block as random factors. In addition, we created a factorial parameter indicating the plasticity of the 2 varieties in the mixture: if both varieties had a positive plasticity in the mixtures, this parameter was “Yes-Yes”; if both varieties had a negative plasticity, it was “No-No”; if one variety had a positive plasticity while the other one had a negative, it was “Yes-No”. This pairwise plasticity response was created for ear-density, and its effect on overyielding, CE, and SE was assessed using linear mixed models with year, site and block as random factors. Pairwise posthoc comparisons were performed with a Tukey test.

#### Relative contributions of CE and SE to overyielding

The relative contributions of CE and SE to overyielding were explored using linear regression models with CE and SE as factors. Effect sizes were calculated as partial R-squares for CE and SE, and reflect the relative contribution of complementarity and selection effects to the yield benefits.

#### Structural Equation Modelling

To investigate the links between plasticity in traits and yield/protein benefits in more details, we applied structural equation modelling (SEM) to our data set. SEM allows to test complex priorly-defined indirect and direct relationships in a single framework and to assess the overall fit of the data to the model (Grace, 2006). The SEMs were built, run, and evaluated with the *lavaan* package in R (Rosseel, 2012). Our *a priori* model was built to relate variety-level plasticity in ear density, biomass, TKW, protein, yield, and plot-level overyielding and overperformance in protein content (equation 1). Plasticity in ear density of both varieties constituted the initial independent variables, because biomass and yield were measured at harvest only, and therefore it is likely that ear density was set priorly to biomass at harvest. Overyielding and overperformance in protein content were the final response variables. Our *a priori* model included the following hypotheses: (1) plasticity in ear density of both varieties directly affects plasticity in biomass, TKW, yield and protein of each variety, (2) plasticity in TKW, yield, and protein of each variety are also affected by plasticity in biomass, (3) plasticity in yield and protein of each variety are affected by plasticity in TKW of the variety, (4) overyielding and overperformance in protein content are affected by the respective plasticity in yield, biomass, and protein of each variety, and (5) all the plasticity values can co-vary with the values from the associated variety.

Because plant height per mixture variety was only assessed in one site (Changins), we created another SEM for Changins only, with height plasticity linked to biomass and ear density plasticity of each variety. All the other traits (TKW, yield, protein content) were subsequently linked to those changes in height in the *a priori* model.

Path coefficients were calculated with maximum likelihood, and the model fits were tested with chi-square goodness of fit test, Bollen-Stine bootstrap test with 1000 bootstrap draws, root mean square error of approximation (RMSEA) test, and comparative fit index (CFI). Non-significant chi-square, Bollen-Stine and RMSEA tests, combined with CFI values above 0.9 indicates a good fit of the model to the data (Kline, 2011; Stefan *et al*., 2021). If the initial model failed the goodness of fit tests, improvements were made with the function *modificationIndices* from the *lavaan* package, which indicates the improvement in model fit (in terms of chi-square statistics) for a selected set of parameters. In addition, matrices of correlations between all variables were computed to identify relevant missing links.

#### Links with phenology

Because our previous analyses indicated a key role of ear density plasticity in overyielding, we investigated this process further. More precisely, we investigated whether the ear density response in mixtures was linked to how plants reacted to light competition during key phenological stages for ear density in wheat. To do so, the link between plasticity in ear density and the phenological response of single plants to shading was assessed with a linear mixed-effects model using *lmer*. For this, we calculated the difference in phenology between the shading treatment and the control treatment, e.g.

> shade-induced phenological plasticity (days) = date of phenological stage in the shade – date of phenological stage in the sun

This phenological plasticity was calculated for the stages related to tillering, i.e. the beginning of tillering (BBCH 21), and the end of tillering (BBCH 29). We chose these two stages as in wheat, ear density is strongly determined by tillering processes (Gawinowski *et al*., 2024). The linear mixed-effects model included tillering-onset plasticity and tillering-end plasticity as fixed factors, and year, site, replicate as well as variety combination as random factors.

## Results

### Plastic responses in mixtures

The varieties did not all respond in the same way when mixed: we observed significant effects of variety on the plasticity of all the measured traits in response to mixtures (Fig. 1, Table 1, Fig. S5, Table S2-S3). Some varieties, such as Bodeli and Campanile, responded positively to being in mixtures, irrespective of the associated variety. They showed increased ear density, biomass and yield in mixtures compared with what was expected from their respective pure stands. In parallel, these varieties had a reduced protein content in mixtures compared to pure stands. The ordination plot further shows that CH211.14074 responded in the same way as Bodeli, and that Molinera responded in a similar way to Campanile. Campanile and Molinera also had higher TKWs in mixtures than in pure stands, which indicates that the increased ear density did not necessarily lead to more but smaller grains. Similarly, Molinera and Bodeli had a higher harvest index in mixtures, suggesting that the increased ear density did not cause more biomass accumulation in stems and leaves at the expense of grain production. On the other side of the ordination plot, we observed CH111.16373 and Colmetta, which both had lower ear density, biomass and yield in mixtures than what was expected from their pure stands, together with a higher protein content. These varieties – along with Schilthorn, whose plastic responses were similar to Colmetta – generally responded negatively to being mixed, irrespective of the associated variety in the mixture. Out of the 8 varieties considered in our study, only Falotta did not show any change in any of the measured traits when grown in mixture.

**Figure 1:**
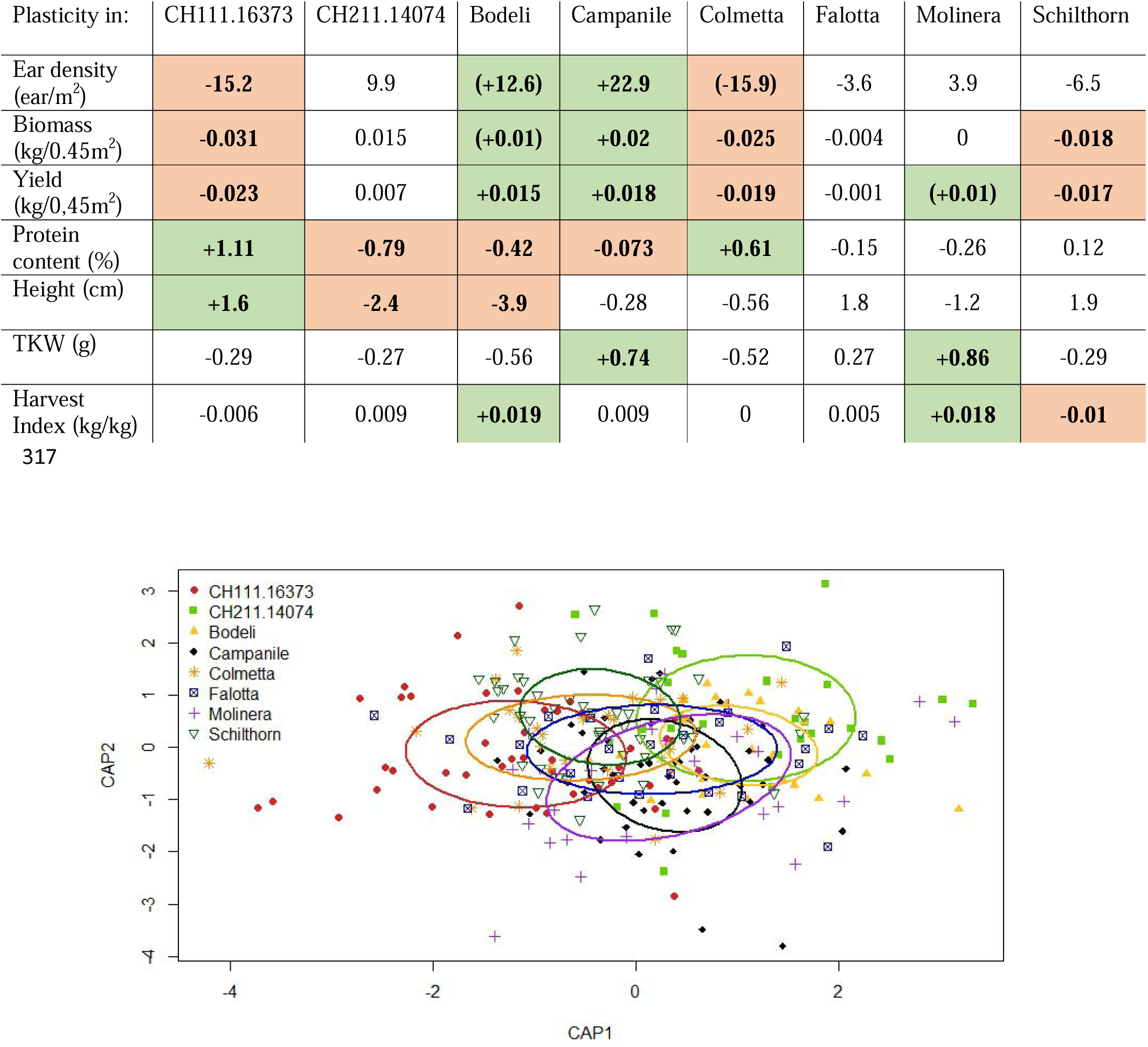
Constrained ordination plot showing changes in mixture-induced trait plasticity in response to variety. Trait plasticity values were averaged per variety × variety combination × site × year. The circles represent the standard deviation of point scores of the ordination. n = 269.

**Table 1:**
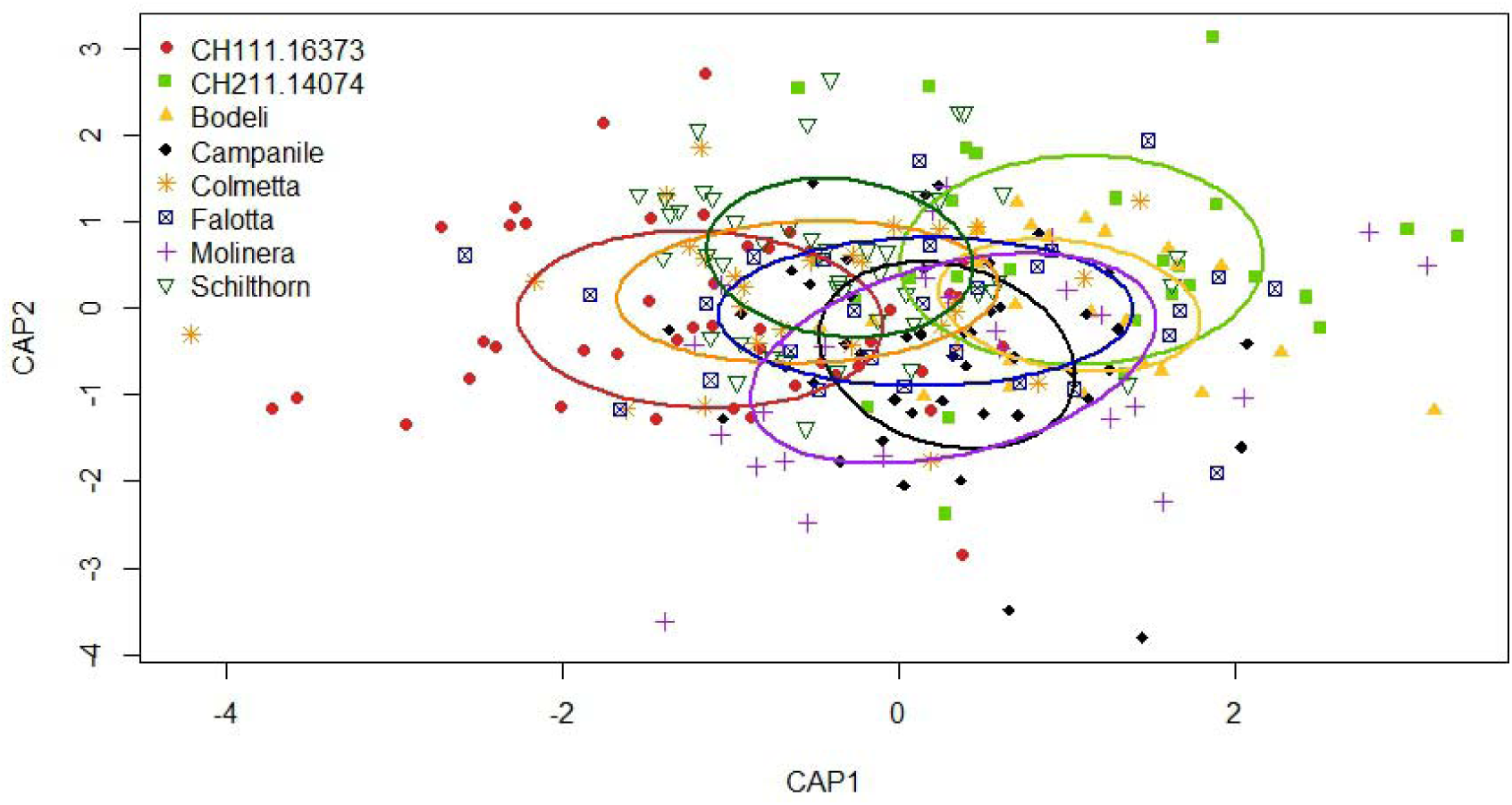
Plasticity of traits in mixtures compared to pure stand. Numbers indicate the mean value of plastic changes averaged per variety × variety combination × site × year. Green cases indicate that the trait was significantly higher in mixtures compared to the corresponding pure stand. Orange cases indicate that the trait value was significantly lower in mixtures compared to pure stand. Significance was defined at α = 0.05 (see Tables S2-S3, Fig S5). Numbers in parentheses indicate p-values between 0.05 and 0.1.

When looking at the specific response of each variety according to the identity of its mixture partner, we found results consistent with the analyses across mixture combinations (Fig. S6, Table S5): Campanile and Bodeli had positive plasticity for yield when mixed with Colmetta/Falotta and CH111.16373/Schilthorn, and positive plasticity for ear density with CH211.14074, Falotta and Schilthorn. CH111.16373 and Schilthorn on the contrary had negative plasticity for biomass and yield when mixed with either Bodeli, Falotta, Molinera or Colmetta. CH111.16373 had a positive plasticity for protein content with most varieties. Results also indicated that most interactions for biomass, yield, and protein content were symmetrical (Fig. S6): this means that in the mixture (A,B), when variety A had an increased biomass or yield, variety B had a decreased biomass or yield.

### Overyielding, Complementarity Effects (CE) and Selection Effects (SE)

Overyielding in absolute value ranged from -0.12 to 0.14 kg/0.45m^2^ (equivalent to -2.6 ton/ha to +3.1 ton/ha), with an average -0.005 kg/0.45m^2^ (i.e. -0.1 ton/ha). Half of the plots had an overyielding between -0.03 and 0.02 kg/0.45m^2^, which translates into -0.6 ton/ha to +0.45 ton/ha. Complementarity effects (CE) and Selection effects (SE) had an average close to 0, with CE in ranges from -0.18 to

0.15 kg/0.45m^2^, and SE from -0.08 to 0.02 kg/0.45m^2^. In other words, the performance of variety mixtures differed more in terms of complementarity than selection effects, meaning that the consequences of changes in plant-plant interactions were more important than the inclusion of a particularly productive variety in the mixture. This is further shown by the relative contributions CE and SE to overyielding: complementarity effects had a significant relative contribution to overyielding and accounted for more than 90% of the variation in overyielding response (r^2^ = 0.903, p-value < 0.0001, Fig. S7). Selection effect did not significantly contribute to overyielding (r^2^ < 0.001, p-value = 0.98). Overyielding was therefore largely driven by complementarity effects.

Overyielding and complementarity effects were significantly reduced by the presence of the variety Colmetta (Fig. 2, Table S7), which notably had a negative biomass and yield plasticity in mixtures. Conversely, including Campanile, which showed positive ear-density, biomass and yield plasticity in mixtures, had a small positive effect on overyielding. SE was significantly reduced by variety combination (Table S6), with Colmetta&Campanile and Colmetta&Schilthorn having a significant lower SE.

**Figure 2:**
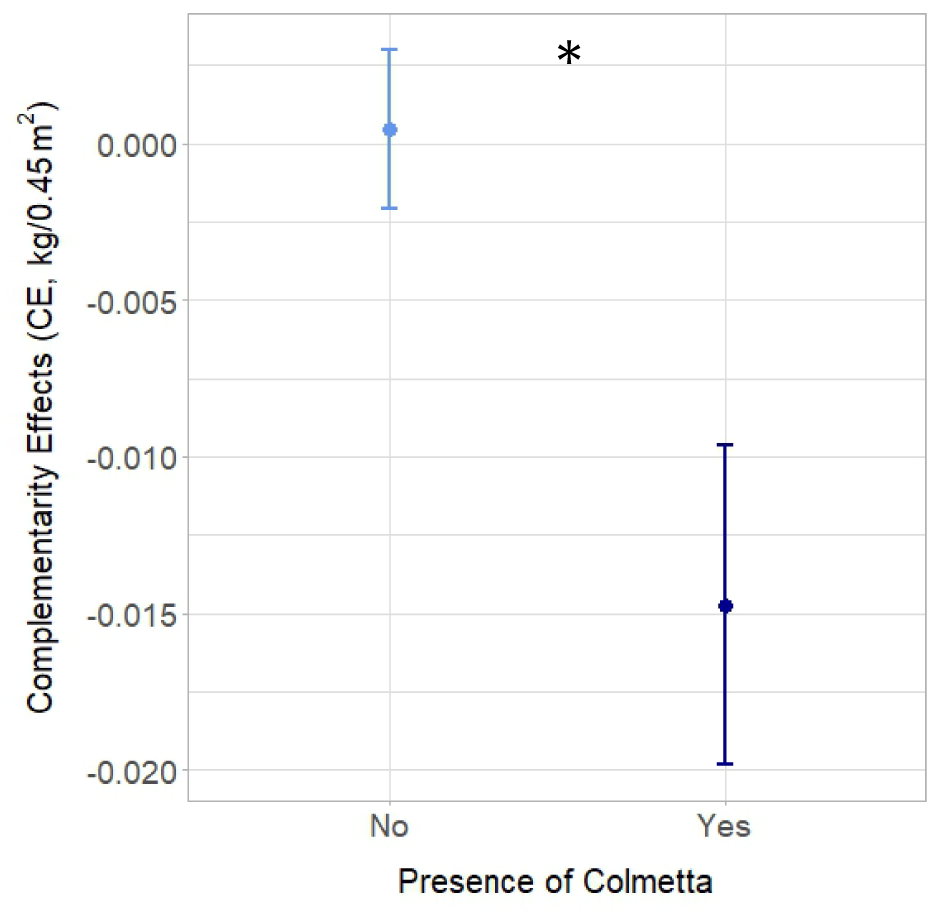
Complementarity effects (CE, kg/0.45m^2^) in response to the absence or presence of the variety Colmetta in the mixture. Dots represent the mean values across replicates, sites, years, and mixture combinations; lines represent the standard error. Stars placed above or next to the results represent the significance of Colmetta presence/absence. n = 401.

Finally, overyielding and CE responded significantly to the pairwise response parameter for ear-density plasticity (Fig. 3, Table S8): they were both significantly higher in Yes-Yes combinations (i.e. when both varieties constituting the mixture had a positive ear density plasticity) and in Yes-No combinations (i.e. when one variety had a positive ear density plasticity and the other one was negative) compared to No-No combinations.

**Figure 3:**
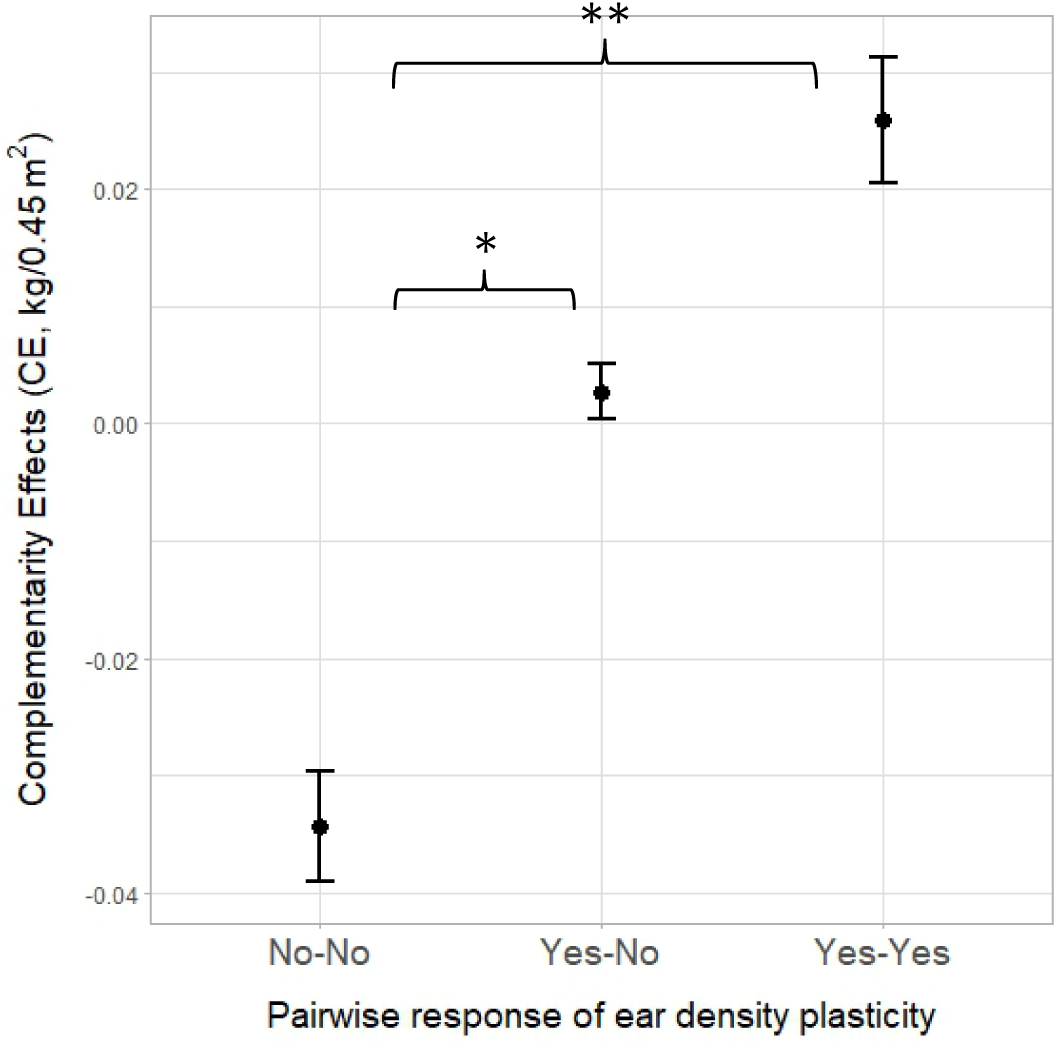
Complementarity Effects (CE) in response to the pairwise parameter for ear density plasticity. Yes-Yes indicates a mixture where both varieties increased their ear densities compared to the respective pure stands corrected by relative sowing densities. Yes-No refers to mixtures where one variety increased its ear density while the second one decreased. No-No indicates mixtures where both varieties decreased their ear densities. n = 401. p-values of the Tukey test: 0.0038 between Yes-Yes and No-No; 0.0126 between Yes-No and No-No

### Structural Equation Modelling

Overall, there was a good fit between our data and the *a priori* SEM: we obtained a chi-square of 22.739, associated with a p-value of 0.121; the p-value of the Bollen-Stine was 0.526; Root Mean Square Error of Approximation (RMSEA) was 0.035 and the associated p-value 0.764; Comparative Fit Index was 0.998. For the second SEM in Changins only, including height plasticity, we obtained a chi-square of 19.7 with a p-value of 0.14; a Bollen-Stine p-value of 0.493; RMSEA was 0.061 with a p-value of 0.346; CFI was 0.995.

In both SEMs, yield benefits were largely driven by positive yield plasticity of the two varieties composing the mixture (Fig. 4, path coefficient of 1.1 between overyielding and yield plasticity for the awned variety, path coefficient of 1.15 between overyielding and yield plasticity for the awnless variety). The means that the yield benefits at the plot-scale were more important when the two varieties had a higher relative yield in the mixtures compared to their respective pure stands. For each variety in the mixture, this plasticity in yield was positively influenced by plasticity in ear density in three ways: once directly via a direct positive correlation, once indirectly via biomass plasticity (which was positively correlated to both yield plasticity and ear-density plasticity) and once even more indirectly via TKW plasticity (positively correlated to yield plasticity and biomass plasticity). This shows that variety-level yield increases in mixtures are largely due to variety-level ear-density increases in mixtures compared to pure stands. Ear-density plasticity of each variety also had a negative impact on yield plasticity of the other variety, but these effects were rather small.

**Figure 4:**
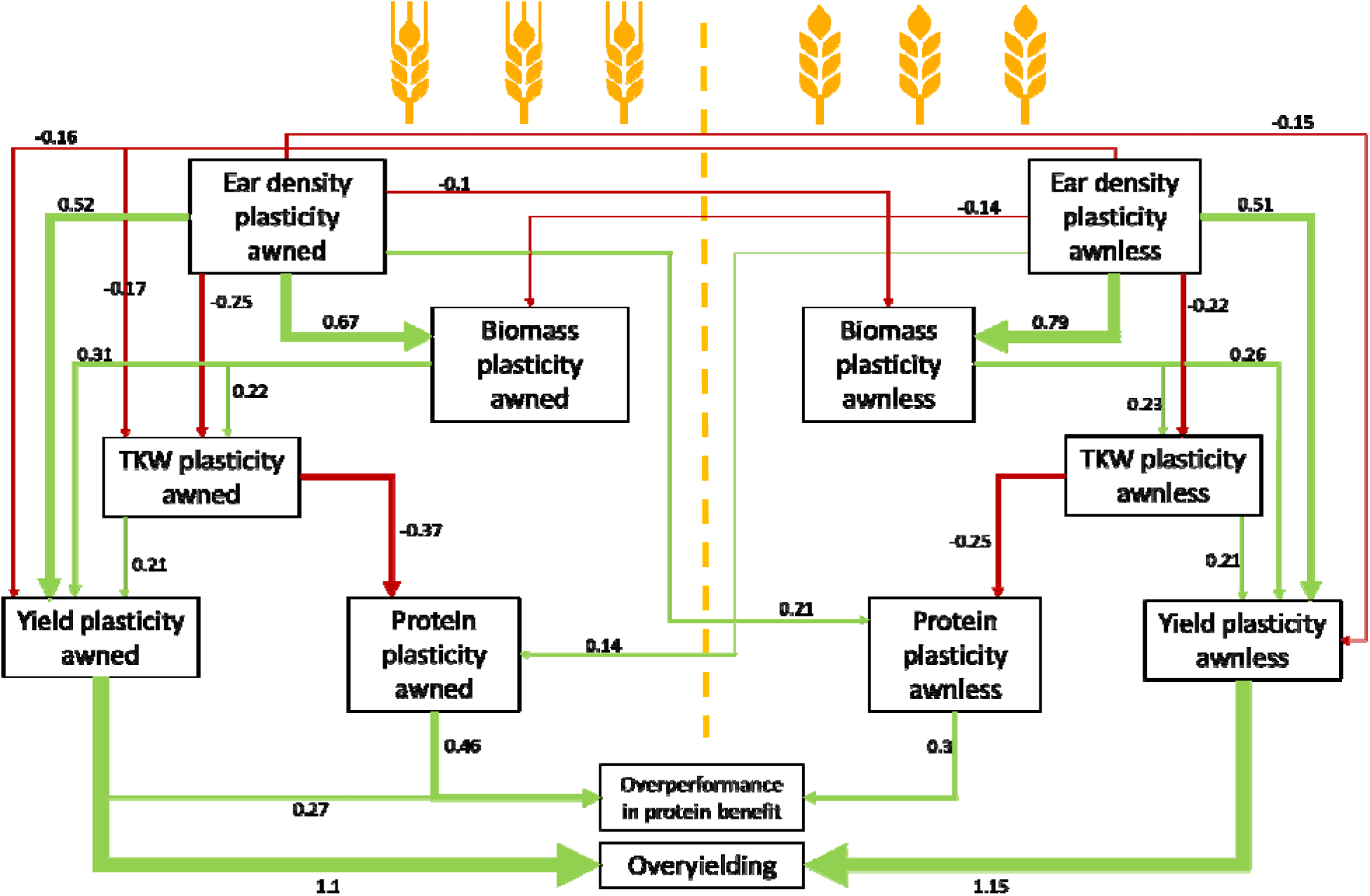
Structural Equation Modeling showing the relationships between plasticity in ear density, biomass, yield, TKW and protein content, and yield and protein benefits (i.e. overyielding and overprotein content). Only significant relationships are shown. Width of the arrows are proportional to the strength of standardized path coefficients indicated by the numbers above/below the arrows (Table S8). Colors of the arrows show positive (green) and negative (red) effects. Residual correlations are not shown. Number of data points: 401.

Ear-density plasticity also positively influences protein benefits, through a negative effect on TKW plasticity, which itself had a negative effect on protein plasticity. Furthermore, the ear-density plasticity of each variety increased protein plasticity of the other variety. Positive protein plasticity from each variety subsequently led to increased protein benefits.

The effects of the awned and awnless varieties on overyielding and overperformance in protein content were mostly symmetrical, in the sense that there was not one mixed variety with higher path coefficients or different pathways than the other mixed variety. The main asymmetrical element was found in the links between yield plasticity and protein benefits, where we saw a positive interaction only for the awned variety.

When looking at the model for Changins only, which included plasticity in height, we found similar links between the variables, notably for plasticity in yield (Fig. S8). In addition, plasticity in height had a direct positive effect on protein plasticity for each mixed variety, indicating that when plants were taller in mixtures, their grains were also more protein-rich. For the awnless varieties, we observed a positive effect of biomass plasticity on height plasticity, but also a negative effect of ear density plasticity on height plasticity, meaning that when there were more ears, the plants were smaller.

### Links with phenology

There was a significant negative correlation between mixture-induced ear density plasticity and shade-induced plasticity for tillering start (Fig. 5). This means that the varieties that had more ears in mixtures compared to pure stands were the one that responded fastest to shading and started tillering earliest under shade. While the correlation was significant, we further observed that 3 varieties did not fit the linear relationship. Colmetta notably fell below the line, meaning that the ear-density plasticity of Colmetta in mixtures was lower than expected from its tillering onset response to shade. Conversely, Campanile and CH211.14074 were above the linear relationship, indicating that their ear-density plasticity in mixtures was higher than expected from their tillering onset response to shade.

**Figure 5:**
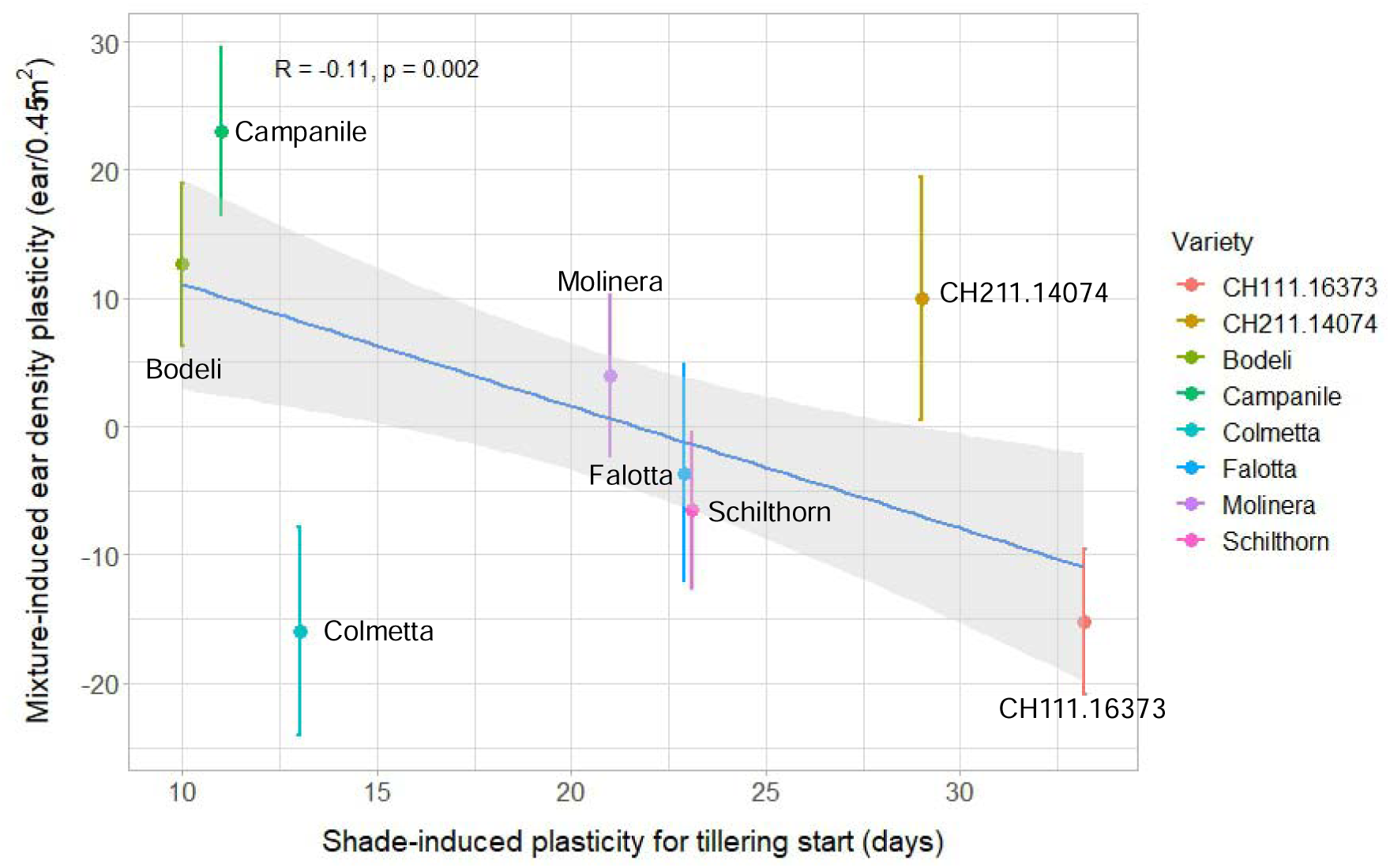
Correlation between ear-density plasticity in mixtures and shade-induced plasticity for tillering-onset (i.e. beginning of tillering in shaded conditions – beginning of tillering in unshaded conditions). Dots represent the mean values per variety, across years, sites, and mixture combinations; lines represent the standard error. Stars placed above represent the significance of the correlation. n = 795.

## Discussion

This study provided a detailed investigation of the processes driving overyielding effects in wheat variety mixtures under realistic agronomic conditions. Our experimental setup and methods allowed us to assess how 8 individual varieties interacted and behaved when grown in 2-variety mixtures compared to pure stands. This was, to our knowledge, one of the few experiments to assess mixture-induced trait plasticity and in real field-like conditions for wheat cultivation, i.e. in replicated field trials involving large plots and no special equipment. Importantly, we were able to measure grain yields of individual varieties in the mixtures, which allowed us to compute not only overyielding, but also its partitioning into complementarity and selection effects *sensu* Hector & Loreau (2001). This is the first time that complementarity and selection effects could be calculated in wheat variety mixtures under realistic conditions, and we could show that yield benefits (i.e. overyielding) were largely dominated by complementarity effects between the varieties. Furthermore, the processes underlying these effects could be partially deciphered thanks to the detailed trait responses of each variety in mixtures, among which we notably pinpointed plasticity in ear density as a key criterion. The additional shading experiment allowed us to dig deeper into the mechanisms of ear density plasticity in mixtures, with the identification of the speed of tillering onset under shade as its main driver.

### Consistency with literature

Trait plasticity in response to mixture environments was previously described in barley cultivar mixtures, where the authors found that each of the 5 studied varieties showed differences in trait responses in mixtures compared to pure stands (Dahlin *et al*., 2020). Similarly, in a study involving 4-cultivar mixtures of wheat, Gawinowski *et al*. found significant differences in height, ear density and tillering dynamics in mixtures vs. pure stands, and these differences were cultivar-specific (Gawinowski *et al*., 2024). Our study likewise emphasized the importance of mixture-induced trait plasticity, with 7 varieties out of 8 showing significant trait changes in mixtures. In our case, some of our wheat varieties – such as Bodeli, Campanile and Molinera – tended to increase their ear numbers and productivity in mixtures while others – Colmetta, CH111.16373 and Schilthorn – were negatively plastic in mixtures for ear density, biomass and yield.

We identified mixture-induced plasticity in ear density as the main driver of overyielding, which was the same conclusion as in the study of Gawinowski (2024). In wheat systems, ear number/density is strongly determined by tillering processes and dynamics (Darwinkel, 1978; Sadras & Slafer, 2012; Slafer *et al*., 2014), which have been proven to be affected by the biotic environment, such as the type of neighbors (Dahlin *et al*., 2020; Blanc *et al*., 2021). In our study we indeed found a strong correlation between ear density plasticity in mixtures and the speed of tillering onset of individual plants under shade, which mimics competition for light (Fig. 5): the more rapidly a variety starts to make tillers in the shade (compared to unshaded conditions), the more fertile tillers it will make in the mixtures. This relationship is coherent, as in a mixed environment, we can expect that the canopy would be more heterogenous, and therefore light availability but also quality can change, which can further affect tillering processes (Casal *et al*., 1986; Evers *et al*., 2006). While plant responses to shading and light competition have been extensively investigated in the field of ecophysiology (Lecarpentier *et al*., 2019; Colbach *et al*., 2020), this is the first time that they are linked to plant plastic changes in mixtures in the field.

With our experiment we could further show that overyielding was largely dominated by complementarity effects between varieties. This is in line with recent results from cultivar experiments in maize, where the authors found that the increase in biomass in mixtures correlated with strong complementarity effects (Su *et al*., 2024). Conversely, selection effects did not play in significant role in driving overyielding. Selection effects arise when a high-performing variety drives the yield of the mixture (Loreau & Hector, 2001; Huang *et al*., 2018). In cultivar mixtures, the differences in yield between the varieties in pure stands are often not large enough to generate strong selection effects in mixtures. In addition, in our experiment the highest yielding variety in pure stands (Colmetta) performed poorly in mixtures and thus tended to decrease mixture yield. With maize cultivar mixtures, Su also identified only neutral or negative selection effects (Su *et al*., 2024).

### Implications for breeders and farmers

Yield components of wheat, as well as their plasticity across environments, have been largely studied for almost a century (Sadras & Slafer, 2012; Sadras & Rebetzke, 2013). However, the bulk of these studies considered wheat as growing in pure stands, and identified key traits and criteria for high performance in monoculture cultivation (Sirat, 2023). Here we show that key traits for high yield in pure stands are different from key traits for high yield in mixtures: for instance, the varieties with the highest ear density in pure stands were not necessarily the ones that had a high relative ear density in mixtures (Fig. S9). If we look at Colmetta, which had a high baseline ear density in pure stands but a negative ear density plasticity in mixture, we can conclude that Colmetta generally suffered in mixtures and was not a good mixing partner. On the contrary, Campanile and Bodeli both had a low baseline ear density in pure stands, but responded positively in mixtures: these two varieties therefore really benefitted from being in a mixture. Similarly, the varieties that exhibited the highest environmental plasticity in monocultures (in response to varying years and sites) were also not necessarily the ones that responded the most to being in mixtures (Fig. S710). Hence, the behaviour of a variety in a mixture cannot be inferred from its behaviour in pure stands; this discrepancy highlights the need to develop new methods and protocols to select and breed varieties for mixtures. The idea that varieties bred for monocultures are not necessarily the best-performing in mixtures has also been examined in intercrop systems, where studies have shown that using mixture-adapted seeds led to higher yield and biomass production than mixing standard, monoculture-adapted germplasm (Stefan *et al*., 2022; Muñoz-Escribano *et al*., 2025).

Yet testing and assessing variety-specific traits in mixtures is tedious and cannot be done for a large number of combinations. This is why is it important to link such behavioural changes in mixtures to intrinsic plant parameters that can be measured more easily. In that sense, the key trait that we identified in the additional shading experiment – i.e. response of tillering onset under shade – is particularly relevant, as it provides an easily measurable parameter on individual plants. This criterion could thus be directly used by breeders to select for good mixture partners, and by farmers and agronomists to create smart mixtures from already existing varieties. For instance, mixing Campanile and Bodeli is more likely to be beneficial for yield, as these two varieties had rapid tillering responses under shade and positive plastic responses for ear density (Fig. S11). On the contrary, mixtures involving Schilthorn or CH111.16373 are not recommendable, as these mixtures would most likely underperform.

### Future research perspectives

The response of tillering onset under shade correlated with ear density plasticity in mixtures for only 5 varieties out of the 8 considered; 3 of the varieties did not fall within the line (Fig. 5). This suggests that, even though response to competition for light clearly plays a major role in ear density plasticity, other mechanisms might be simultaneously at play in mixtures. For instance, here we only recorded the timing of tillering processes, but the number of fertile tillers per plant is ultimately determined by tiller emission and regression, the latter depending on both competition for light and for nitrogen (Ding *et al*., 2021; Gawinowski *et al*., 2024). A more detailed investigation into tillering dynamics, including following how many tillers are created and how many regress, would be helpful to dig deeper into this. However, these measurements are cumbersome and would greatly benefit from digital phenotyping methods (David *et al*., 2023). In addition to this, belowground processes – which were not considered in this study – have been shown to influence plant-plant interactions in diverse systems, either via plasticity and changes in root traits (Montazeaud *et al*., 2018; Colom & Baucom, 2020), changes in microbiota (Latz *et al*., 2015; Stefan *et al*., 2021), or chemical signalling (Zhang *et al*., 2016; Hierro & Callaway, 2021). Most importantly, recent work has shown that plants can recognize the identity of their neighbour (i.e. kin recognition) and distinguish whether the neighbours are the same genotype or not (Dudley & File, 2007; Biedrzycki *et al*., 2011; Fréville *et al*., 2019). The mechanisms of this kin recognition remain incomplete; some evidence points towards the role of light patterns received by the plants (Crepy & Casal, 2015), but most research emphasizes the role of root exudates (Semchenko *et al*., 2014; Depuydt, 2014). Metabolites expressed by the roots can indeed carry specific information regarding genetic relatedness of the neighbours, and can then induce different responses at the plant level (Fang *et al*., 2013; Monchgesang *et al*., 2016). For instance, the presence and recognition of intraspecific neighbours through soil-located processes can modify plant immunity and disease susceptibility in wheat and rice (Subrahmaniam *et al*., 2018; Pélissier *et al*., 2021, 2023). Further research is thus needed to understand the specific behaviour of our varieties that responded more or less than expected to the mixture environment based on their tillering onset under shade. Finding the molecular or even genetic mechanisms would help breeding programs to further promote the selection of good mixing partners (Wuest *et al*., 2021; Becker *et al*., 2023).

Furthermore, our results indicated that most interactions in mixtures were symmetrical: we observed compensative mechanisms (Fig. S6) between the varieties, with one variety increasing its relative ear density or yield while the other one was decreasing its relative ear density or yield. Thus, the key is to find partners that either both manage to increase their relative yield/ear density in mixtures (positive-positive), or to find one partner increasing its relative yield/ear density while the second is staying stable (positive-neutral). This happened for a few combinations in our experiment: for instance, in the mixture Campanile&CH211.14074, Campanile increased its relative ear density while CH211.14074 stayed the same. Campanile and CH211.14074 are the two varieties that performed better in mixtures than expected based on their tillering phenology response under shade (Fig. 5); interestingly, these two varieties are both spring type with no or very low vernalization requirement. It would thus be interesting to test other varieties close to spring wheat to see whether they also make good mixture partners. The pairwise characteristic of plant-plant interactions has been emphasized in a large number of studies investigating inter- and intraspecific mixtures (Newton *et al*., 2008; Dahlin *et al*., 2020; Chen *et al*., 2021; Cossani & Sadras, 2021). These studies showed that the behaviour of varieties/species in mixtures was generally dependent on the identity of the neighbour, which makes inferences about good mixture partners even more complex (Dahlin *et al*., 2020; Chen *et al*., 2021). This is why different approaches based on pairwise responses and network analyses of genes or traits could offer important insights into how to combine varieties/species for optimal productivity (Wuest *et al*., 2023; Lebreton *et al*., 2024; Mathieu *et al*., 2024).

## Conclusions

Using real-conditions field experiments where we assessed variety-specific trait and productivity changes in mixtures, this study assessed, for the first time, the plastic processes driving overyielding, complementarity and selection effects in wheat variety mixtures. Overyielding was largely dominated by complementarity effects, which explained more than 90% of its variance. Furthermore, we identified mixture-induced plasticity in ear density, i.e. the ability of the mixed varieties to increase their ear density in mixtures compared to pure stands, as the main driver of overyielding. An additional experiment using the response of individual plants to shade allowed us to link plastic changes in ear density in mixtures to the timing of tillering onset under shade. Overall, our study emphasizes the importance of considering variety-specific plastic changes in traits in response to the presence of non-kin neighbours when designing variety mixtures. Indeed, varieties are generally selected as mixture partners based on their phenotype in pure stands; here, we demonstrate that what really matters is whether and how these varieties are able to change their phenotype and tillering ability in mixtures. While further research is needed to identify the molecular and genetic bases of this plasticity in response to mixtures, the identification of a new key trait – tillering onset under shade – that is easily measurable on individual plants already opens new possibilities for mixture-targeted breeding programs and for successful mixture combinations.

## Acknowledgements

We thank Yann Imhoff, Noémie Schaad, Julie Roux, Reynold Bovy, Gabriel Dessiex, Flavio Foiada and Malgorzata Watroba for their assistance with field experiments and post-harvest analyses. We also acknowledge the support from Hans Winzeler regarding the choice of the accessions, and the design of the experimental design. This project was jointly funded by the Swiss Federal Office for Agriculture (BLW) and IP-Suisse.

## Author contributions

**Laura Stefan:** Investigation, Data Curation, Formal analysis, Project administration, Visualization, Writing – Original Draft, Writing – Review and Editing. **Silvan Strebel:** Investigation, Writing – Review and Editing. **Nathalie Colbach:** Conceptualization, Funding acquisition, Resources, Writing – Review and Editing. **Dario Fossati:** Conceptualization, Funding acquisition, Resources, Writing – Review and Editing. **Lilia Levy Häner:** Conceptualization, Funding acquisition, Resources, Investigation, Supervision, Writing – Review and Editing.

## Data availability

Data is available on Zenodo: https://doi.org/10.5281/zenodo.14917728.

## Supporting information

Supplementary data associated with this article can be found online.

