## Supplementary Material for "Plasticity in ear density drives complementarity effects and yield benefits in wheat variety mixtures"

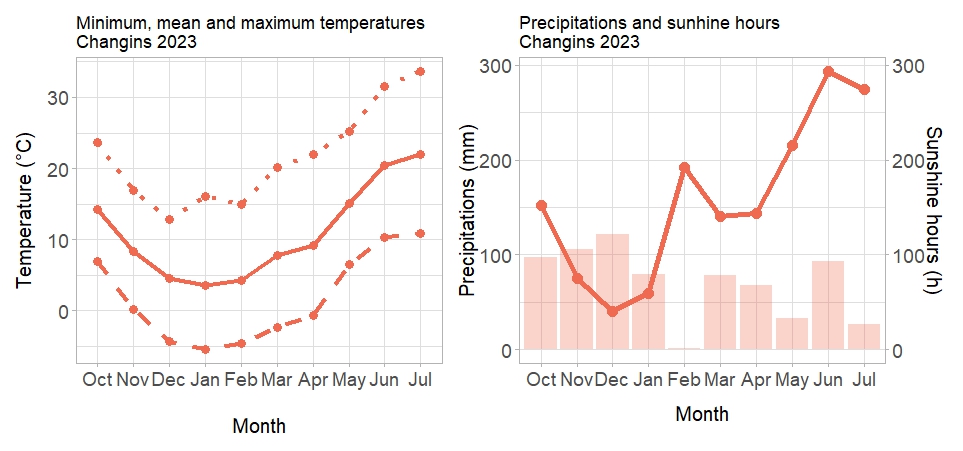

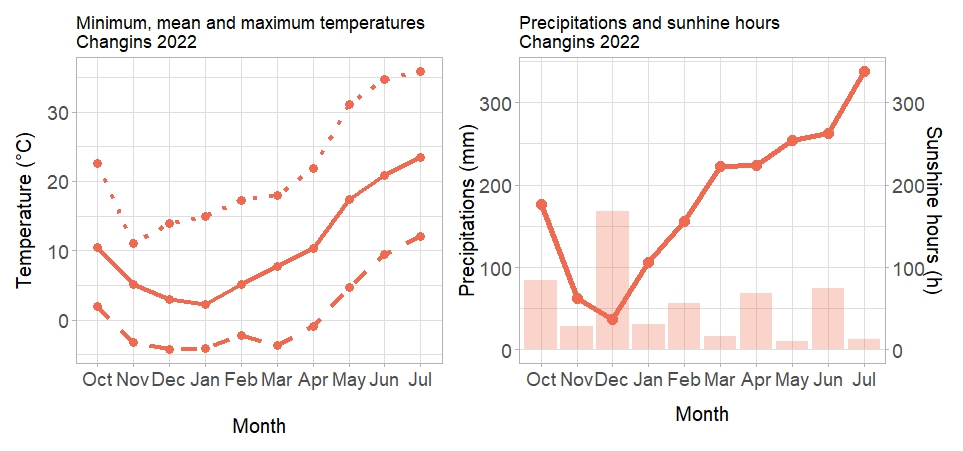

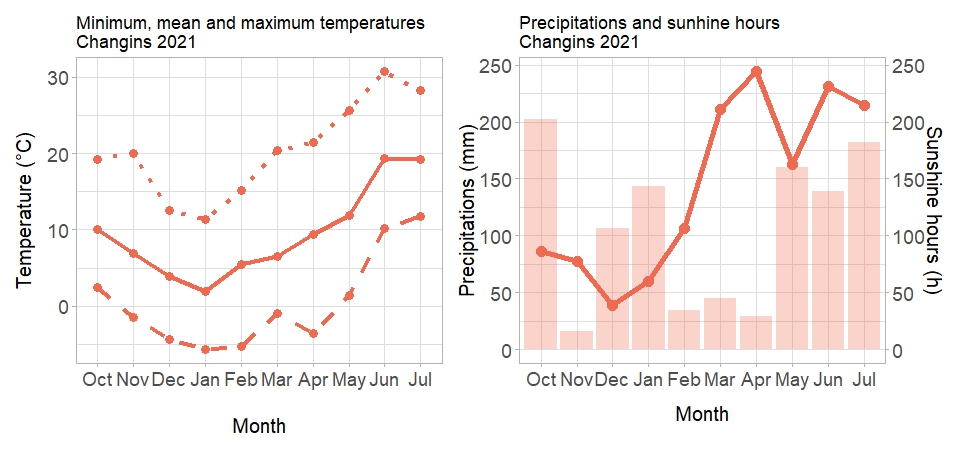

**Fig. S1: Left panel: Minimum, mean, and maximum temperatures in Changins 2021 (a), 2022 (b) and 2023 (c).** Dashed lines represent minimum temperatures, full lines represent the mean, and dotted lines represent maximum temperatures. **Right panel: Precipitations and sunshine hours.** Lines represent the sunshine hours, while bars show the precipitation.

c

**(a)**

c

**(b)**

c

**(c)**

c

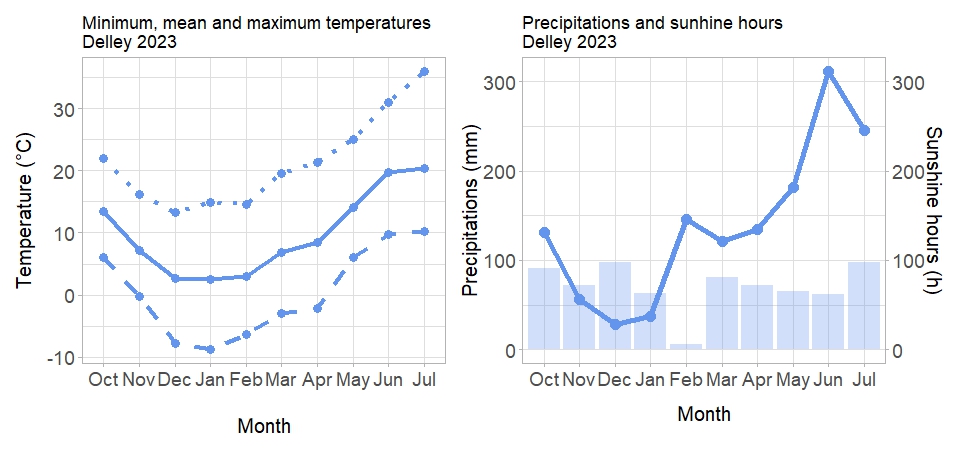

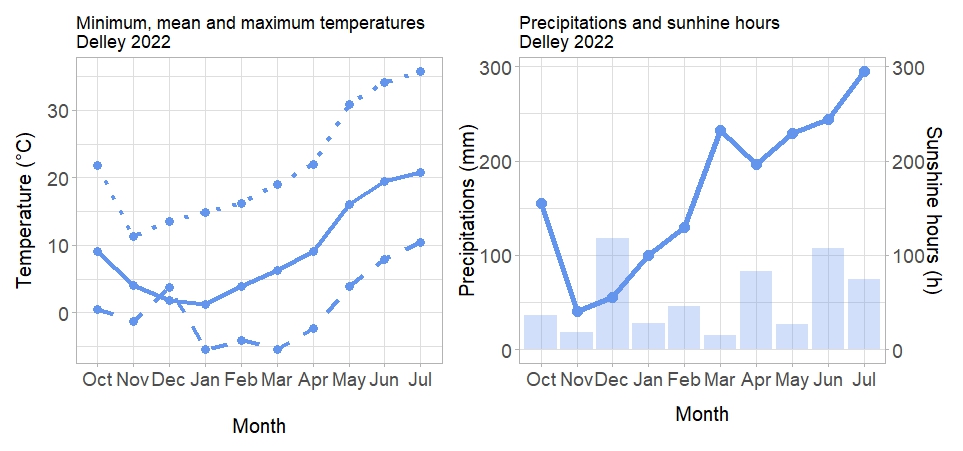

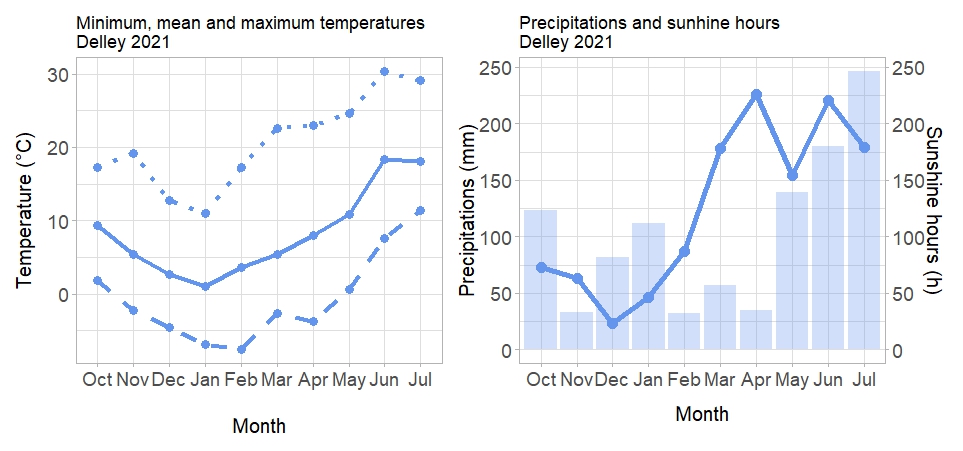

**Fig. S2: Left panel: Minimum, mean, and maximum temperatures in Delley 2021 (a), 2022 (b) and 2023 (c).** Dashed lines represent minimum temperatures, full lines represent the mean, and dotted lines represent maximum temperatures. **Right panel: Precipitations and sunshine hours.** Lines represent the sunshine hours, while bars show the precipitation.

c

**(a)**

c

**(b)**

c

**(c)**

c

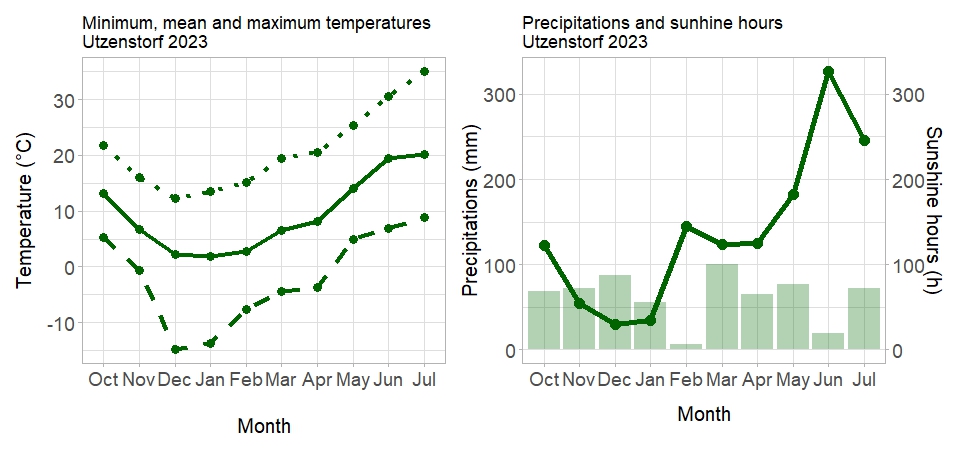

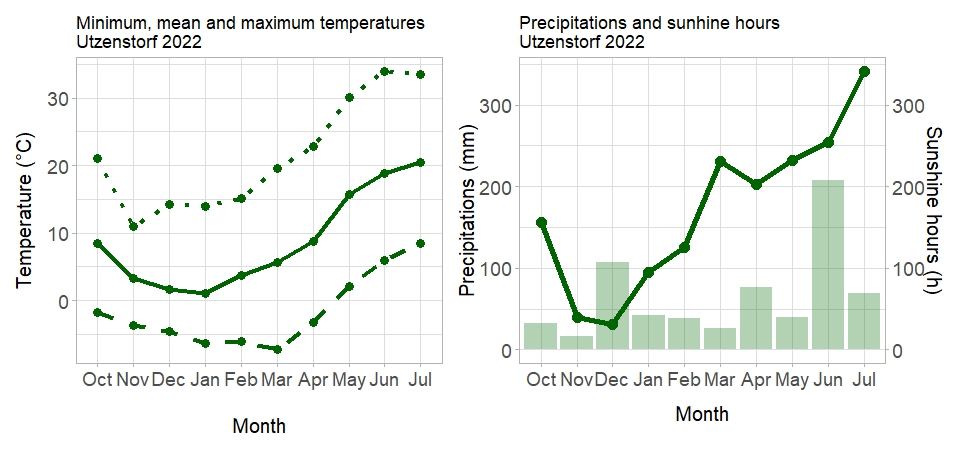

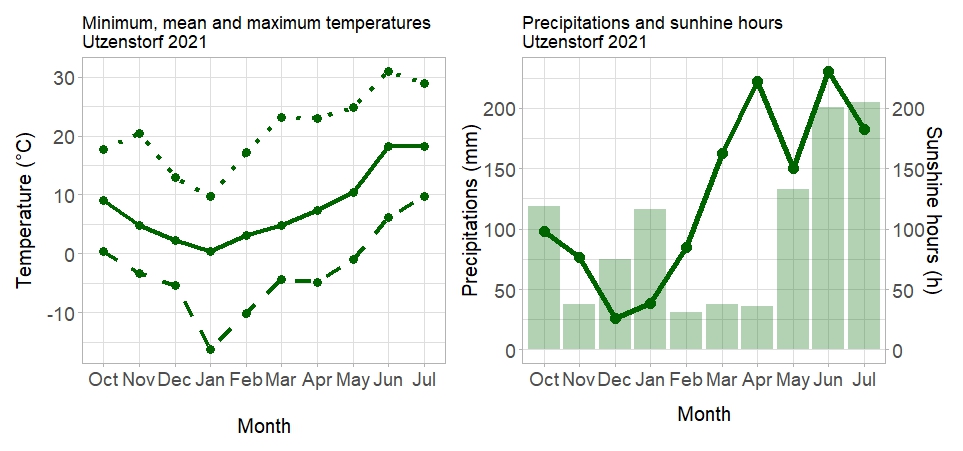

**Fig. S3: Left panel: Minimum, mean, and maximum temperatures in Utzenstorf 2021 (a), 2022 (b) and 2023 (c).** Dashed lines represent minimum temperatures, full lines represent the mean, and dotted lines represent maximum temperatures. **Right panel: Precipitations and sunshine hours.** Lines represent the sunshine hours, while bars show the precipitation.

c

**(a)**

c

**(b)**

c

**(c)**

c

**Figure S4:** Schematic illustration of the method to sort the components of the mixtures and of the post-harvest analyses.

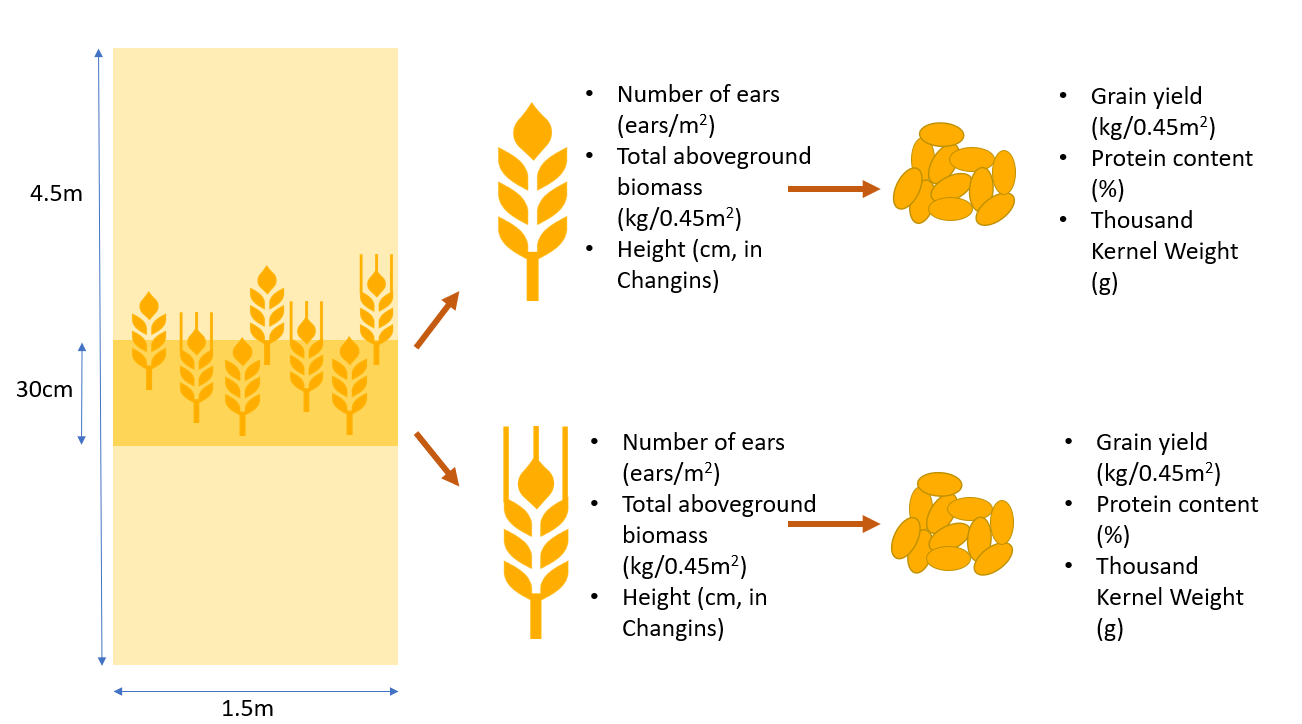

**Table S1:** Experimental details and soil status of the fields used for the experimental trials

|  | *Changins 2021* | *Changins 2022* | *Changins 2023* | *Delley 2021* | *Delley 2022* | *Delley 2023* | *Utzenstorf 2021* | *Utzenstorf 2022* | *Utzenstorf 2023* |
| --- | --- | --- | --- | --- | --- | --- | --- | --- | --- |
| *Sowing date* | 20.10.2020 | 13.10.2021 | 18.10.2022 | 16.10.2020 | 15.10.2021 | 18.10.2022 | 20.10.2020 | 28.10.2021 | 19.10.2022 |
| *Harvest date* | 23.07.2021 | 11.07.2022 | 11.07.2023 | 23.07.2021 | 13.07.2022 | 11.07.2023 | 23.07.2021 | 18.07.2022 | 17.07.2023 |
| *% Clay* | 26 | 22 | 21 | 14 | 20 | 15 | 15 – 20 | 15 – 20 | 15 – 20 |
| *% Silt* | 43 | 47 | 43 | 28 | 46 | NA | NA | NA | NA |
| *% Sand* | 31 | 31 | 36 | 58 | 34 | NA | NA | NA | NA |
| *% Organic Matter* | 2.9 | 2.5 | 2.2 | 1.5 | 2.2 | 1.5 | 2 – 4.9 | 2 – 4.9 | 2 – 4.9 |
| *pH* | 7.7 | 7.6 | 7.1 | 6.9 | 8 | 6.3 | 6.4 | 5.7 | 6.9 |

**Table S2**: Type-III Analysis of Variance Table of the different trait plasticity responses to variety.

*SumSQ,* sum of squares*, MeanSQ,* mean square of error*, DenDF*, degrees of freedom of error term; *NumDF*, degrees of freedom of term; *F-value*, variance ratio; p-values in bold are signiﬁcant at α = 0.05; * (P < 0.05), ** (P < 0.01), *** (P < 0.001). n = 795 for all the traits except height (n = 264).

|  | *Sum SQ* | *Mean SQ* | *NumDF* | *DenDF* | *F-value* | *p-value* |
| --- | --- | --- | --- | --- | --- | --- |
| *Ear density* | 188138 | 23517 | 8 | 42.948 | 4.9 | < 0.001 *** |
| *Biomass* | 0.38 | 0.047 | 8 | 56.8 | 9.7 | < 0.001 *** |
| *Yield* | 0.29 | 0.036 | 8 | 67.7 | 20.4 | < 0.001 *** |
| *Height* | 945.15 | 118.14 | 8 | 13.148 | 5.14 | 0.0046 ** |
| *Protein* | 394.14 | 49.3 | 8 | 100.8 | 26.57 | < 0.001 *** |
| *Harvest Index* | 0.08 | 0.01 | 8 | 23.9 | 5.1 | < 0.001 *** |
| *TKW* | 207 | 25.88 | 8 | 28.37 | 3.54 | 0.0058 ** |

**Table S3:** Variety-level significance derived from the Analysis of variance above

| **Ear density** | Estimate | Std. Error | df | t value | Pr(>\|t\|) |  |
| --- | --- | --- | --- | --- | --- | --- |
| *CH111.16373* | -18.99 | 7.93 | 37.76 | -2.40 | **0.02** | ***** |
| *CH211.14074* | 5.07 | 9.86 | 51.00 | 0.51 | 0.61 |  |
| *Bodeli* | 19.02 | 9.91 | 51.45 | 1.92 | **0.06** | **.** |
| *Campanile* | 32.91 | 7.98 | 38.77 | 4.12 | **< 0.001** | ******* |
| *Colmetta* | -18.31 | 9.81 | 50.89 | -1.87 | **0.07** | **.** |
| *Falotta* | -5.26 | 9.85 | 51.66 | -0.53 | 0.60 |  |
| *Molinera* | 6.26 | 9.82 | 50.57 | 0.64 | 0.53 |  |
| *Schilthorn* | -13.10 | 7.91 | 37.54 | -1.66 | 0.11 |  |
| **Biomass** | Estimate | Std. Error | df | t value | Pr(>\|t\|) |  |
| *CH111.16373* | -0.04 | 0.01 | 41.66 | -4.91 | **< 0.001** | ******* |
| *CH211.14074* | 0.01 | 0.01 | 62.69 | 1.17 | 0.25 |  |
| *Bodeli* | 0.02 | 0.01 | 63.86 | 1.91 | **0.06** | **.** |
| *Campanile* | 0.04 | 0.01 | 43.28 | 4.15 | **< 0.001** | ******* |
| *Colmetta* | -0.03 | 0.01 | 65.36 | -3.05 | **0.003** | ****** |
| *Falotta* | 0.00 | 0.01 | 62.88 | -0.43 | 0.67 |  |
| *Molinera* | 0.00 | 0.01 | 63.35 | 0.10 | 0.92 |  |
| *Schilthorn* | -0.02 | 0.01 | 43.97 | -2.65 | **0.01** | ***** |
| **Yield** | Estimate | Std. Error | df | t value | Pr(>\|t\|) |  |
| *CH111.16373* | -0.04 | 0.01 | 36.54 | -6.02 | **< 0.001** | ******* |
| *CH211.14074* | 0.01 | 0.01 | 62.53 | 0.94 | 0.35 |  |
| *Bodeli* | 0.03 | 0.01 | 63.20 | 3.48 | **< 0.001** | ******* |
| *Campanile* | 0.03 | 0.01 | 37.80 | 5.58 | **< 0.001** | ******* |
| *Colmetta* | -0.03 | 0.01 | 62.02 | -3.94 | **< 0.001** | ******* |
| *Falotta* | -0.01 | 0.01 | 62.02 | -0.73 | 0.47 |  |
| *Molinera* | 0.01 | 0.01 | 62.59 | 1.77 | **0.08** | **.** |
| *Schilthorn* | -0.02 | 0.01 | 38.24 | -3.31 | **0.002** | ****** |
| **Height** | Estimate | Std. Error | df | t value | Pr(>\|t\|) |  |
| *CH111.16373* | 2.49 | 1.04 | 14.07 | 2.41 | **0.03** | ***** |
| *CH211.14074* | -3.01 | 1.25 | 25.25 | -2.40 | **0.02** | ***** |
| *Bodeli* | -3.62 | 1.25 | 25.25 | -2.89 | **0.01** | ****** |
| *Campanile* | -0.83 | 1.04 | 14.07 | -0.81 | 0.43 |  |
| *Colmetta* | 0.31 | 1.30 | 28.46 | 0.24 | 0.81 |  |
| *Falotta* | 1.84 | 1.25 | 25.25 | 1.48 | 0.15 |  |
| *Molinera* | -1.91 | 1.25 | 25.25 | -1.53 | 0.14 |  |
| *Schilthorn* | 1.54 | 1.06 | 15.09 | 1.46 | 0.17 |  |
| **Protein** | Estimate | Std. Error | df | t value | Pr(>\|t\|) |  |
| *CH111.16373* | 1.79 | 0.24 | 35.76 | 7.55 | **< 0.001** | ******* |
| *CH211.14074* | -0.93 | 0.27 | 56.75 | -3.42 | **0.001** | ****** |
| *Bodeli* | -0.74 | 0.27 | 57.36 | -2.69 | **0.01** | ****** |
| *Campanile* | -0.57 | 0.24 | 37.09 | -2.36 | **0.02** | ***** |
| *Colmetta* | 1.12 | 0.27 | 55.36 | 4.14 | **< 0.001** | ******* |
| *Falotta* | -0.16 | 0.27 | 55.84 | -0.58 | 0.57 |  |
| *Molinera* | -0.32 | 0.27 | 55.56 | -1.17 | 0.25 |  |
| *Schilthorn* | -0.05 | 0.24 | 35.80 | -0.20 | 0.85 |  |
| **HI** | Estimate | Std. Error | df | t value | Pr(>\|t\|) |  |
| *CH111.16373* | -0.01 | 0.01 | 27.32 | -1.37 | 0.18 |  |
| *CH211.14074* | 0.01 | 0.01 | 34.18 | 1.51 | 0.14 |  |
| *Bodeli* | 0.02 | 0.01 | 32.77 | 3.26 | **0.003** | ****** |
| *Campanile* | 0.01 | 0.01 | 28.96 | 1.96 | 0.06 |  |
| *Colmetta* | 0.00 | 0.01 | 33.57 | -0.05 | 0.96 |  |
| *Falotta* | 0.00 | 0.01 | 31.57 | 0.87 | 0.39 |  |
| *Molinera* | 0.02 | 0.01 | 32.81 | 3.25 | **0.003** | ****** |
| *Schilthorn* | -0.01 | 0.01 | 29.51 | -2.31 | **0.03** | ***** |
| **TKW** | Estimate | Std. Error | df | t value | Pr(>\|t\|) |  |
| *CH111.16373* | -0.42 | 0.29 | 30.41 | -1.43 | 0.16 |  |
| *CH211.14074* | -0.39 | 0.36 | 36.25 | -1.08 | 0.29 |  |
| *Bodeli* | -0.56 | 0.36 | 33.46 | -1.57 | 0.13 |  |
| *Campanile* | 0.83 | 0.29 | 30.20 | 2.84 | **0.01** | ****** |
| *Colmetta* | -0.55 | 0.35 | 33.25 | -1.57 | 0.13 |  |
| *Falotta* | 0.33 | 0.36 | 36.61 | 0.91 | 0.37 |  |
| *Molinera* | 0.88 | 0.36 | 34.94 | 2.46 | **0.02** | ***** |
| *Schilthorn* | -0.27 | 0.29 | 29.85 | -0.92 | 0.36 |  |

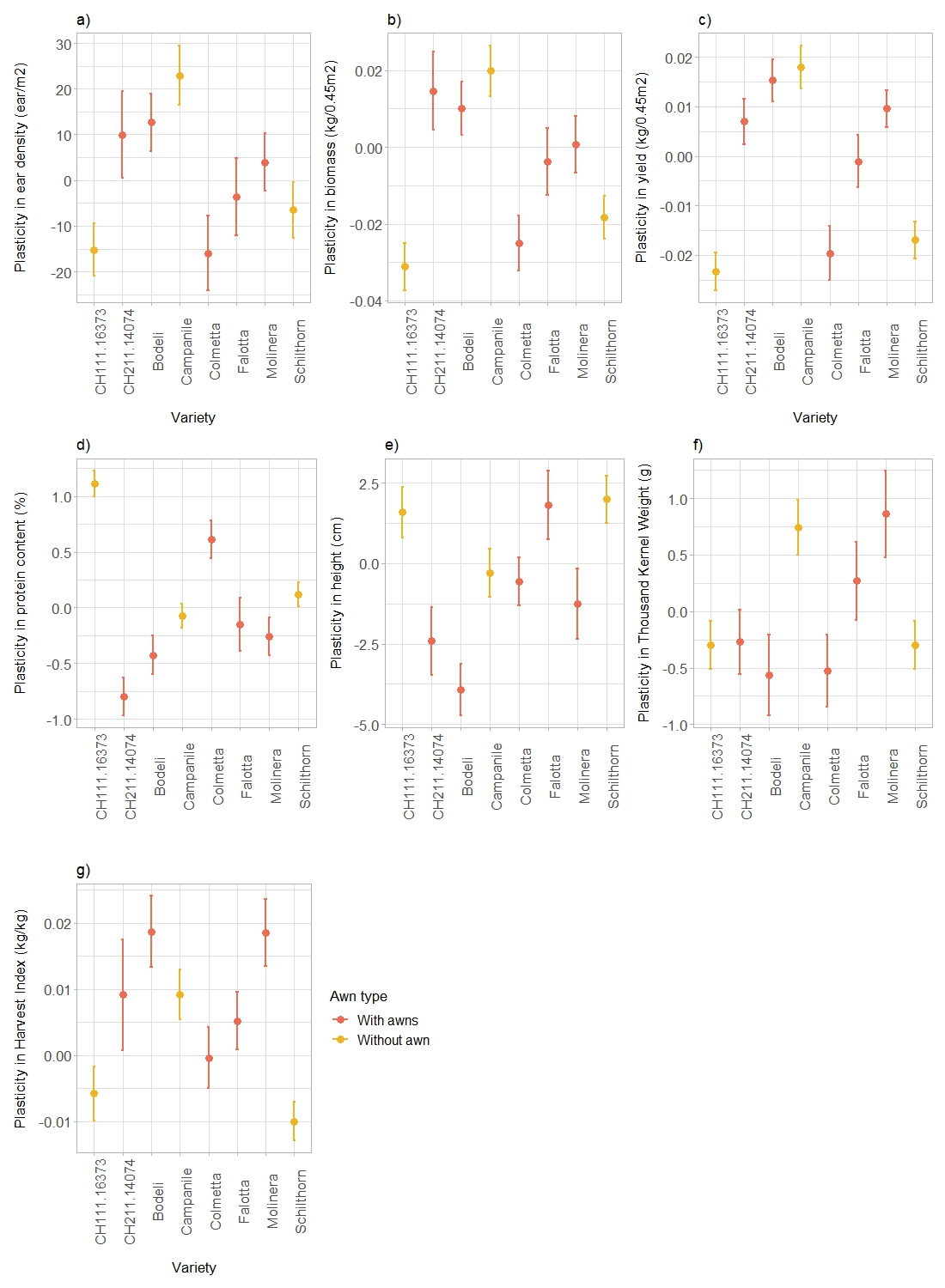

**Figure S5.** Mixture-induced plasticity in ear density (a), biomass (b), yield (c), protein content (d), height (e), TKW (f), and harvest index (g) in mixtures, averaged across sites, years, and mixture combinations. See Table S3 for the statistical details. n = 795 for all the traits except height (n = 264).

**Table S4:** Means of all measured traits across all plots and sites

|  | Ear density (ear/m^2^) | Aboveground biomass (ton/ha) | Grain yield (ton/ha) | Protein content (%) | Height (cm) | Thousand Kernel Weight (g) | Harvest Index (kg/kg) | Tillering onset unshaded (DOY) | Tillering end unshaded (DOY) |
| --- | --- | --- | --- | --- | --- | --- | --- | --- | --- |
| Average across all plots | 502 | 9.8 | 6.4 | 12.6 | 96 | 39 | 0.39 | 334 | 110 |

|  | Density plasticity | | Biomass plasticity | | Yield plasticity | | Protein content plasticity | | Height plasticity | | TKW plasticity | | HI plasticity | |
| --- | --- | --- | --- | --- | --- | --- | --- | --- | --- | --- | --- | --- | --- | --- |
| Focal variety | F value | Pr(F) | F value | Pr(F) | F value | Pr(F) | F value | Pr(F) | F value | Pr(F) | F value | Pr(F) | F value | Pr(F) |
| CH111.16373 | 1.55 | 0.18 | 3.05 | **0.015** | 2.9 | **0.029** | 8.38 | **< 0.001** | 2.13 | 0.1 | 1.49 | 0.22 | 1.6 | 0.17 |
| CH211.14074 | 16.6 | 0.68 | 2.08 | 0.14 | 3.56 | **0.037** | 7.1 | **0.0029** | 3.01 | **0.073** | 2.6 | **0.066** | 1.5 | 0.24 |
| Bodeli | 1.57 | 0.23 | 2.67 | **0.08** | 4.99 | **0.012** | 4.8 | **0.0057** | 6.6 | **0.0073** | 1.66 | 0.19 | 2.3 | 0.11 |
| Campanile | 1.98 | 0.12 | 2.56 | **0.055** | 3.18 | **0.023** | 1.69 | 0.15 | 0.65 | 0.66 | 1.55 | 0.18 | 1.17 | 0.35 |
| Colmetta | 6.01 | **0.0057** | 9.2 | **< 0.001** | 11.41 | **< 0.001** | 16 | **< 0.001** | 0.61 | 0.67 | 2.19 | 0.1 | 0.98 | 0.42 |
| Falotta | 2.8 | **0.07** | 3.9 | **0.015** | 7.1 | **< 0.001** | 9.4 | **< 0.001** | 3.23 | 0.24 | 3.39 | **0.043** | 2.28 | **0.09** |
| Molinera | 3.94 | **0.027** | 4.04 | **0.025** | 11.02 | **< 0.001** | 7.8 | **0.001** | 7.9 | 0.11 | 0.97 | 0.43 | 5.05 | **0.011** |
| Schilthorn | 1.28 | 0.29 | 2.69 | **0.043** | 2.90 | **0.029** | 4.16 | **0.0062** | 1.75 | 0.38 | 1.27 | 0.3 | 1.49 | 0.23 |

**Table S5:** Type-III Analysis of Variance Table of the trait plasticity responses of each variety (i.e. focal variety) to the identity of the mixture partner. The first column indicates the focal variety that is investigated. The subsequent columns indicate the response to mixture partner identity for the plasticity of each of the traits. TKW: Thousand Kernel Weight; HI: Harvest Index.

*F-value*, variance ratio; *Pr(>F)*, error probability. P-values in bold are signiﬁcant at α = 0.05; * (P < 0.05), ** (P < 0.01), *** (P < 0.001).

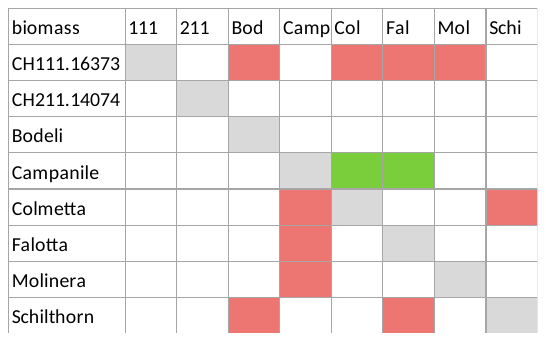

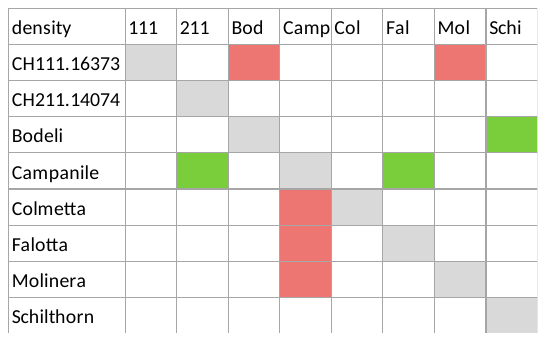

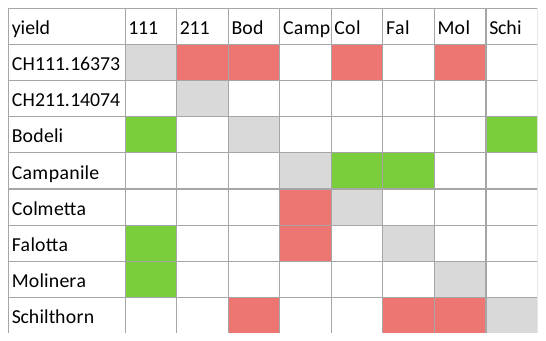

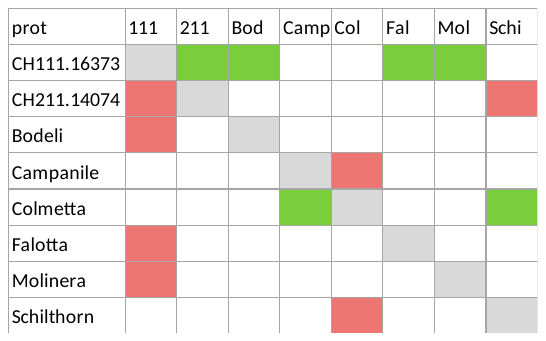

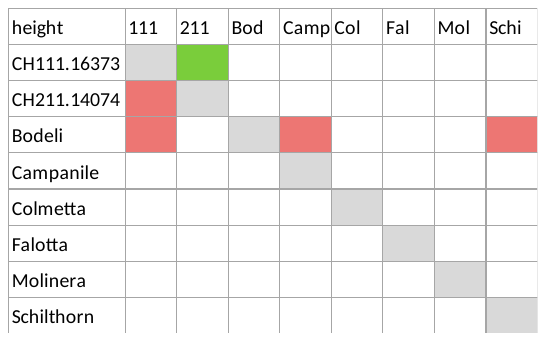

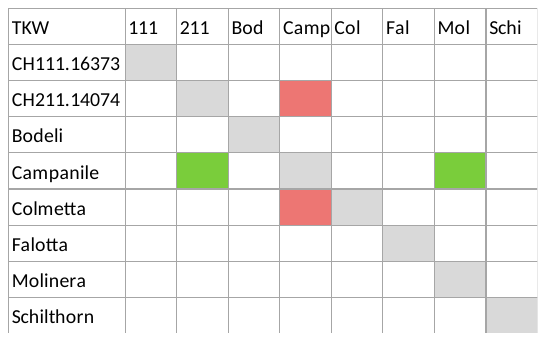

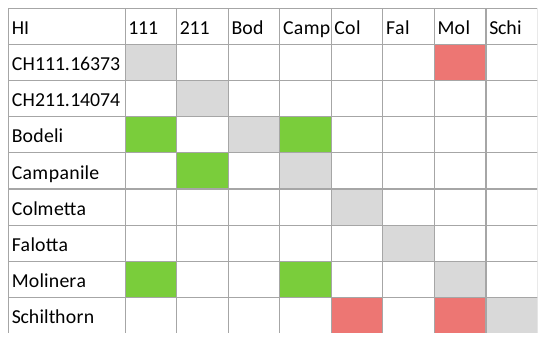

**Figure S6.** Variety-specific responses of trait plasticity to the identity of the mixture partner: the first column indicates the focal variety, while the first row indicates the identity of the mixture partner. Green cases indicate a significant positive plastic response, while red cases indicate a significant negative plastic response. Example: for ear density plasticity, CH111.16373 had a negative plastic response (i.e. lower ear density compared to pure stand corrected by relative sowing density) when mixed with Bodeli or Molinera. Campanile had a positive response (i.e. higher ear density compared to pure stand corrected by relative sowing density) when mixed with CH211 or Falotta.

Abbreviations: Density: ear density; TKW: Thousand Kernel Weight; HI: Harvest Index; prot: protein content.

**Table S6.** Type-III Analysis of Variance Table of overyielding, Complementarity Effects (CE) and Selection Effects (SE), and overperformance in protein content in response to variety combination.

*SumSQ,* sum of squares*, MeanSQ,* mean square of error*, DenDF*, degrees of freedom of error term; *NumDF*, degrees of freedom of term; *F-value*, variance ratio; p-values in bold are signiﬁcant at α = 0.05; * (P < 0.05), ** (P < 0.01), *** (P < 0.001). n = 401.

|  | Sum Sq | Mean Sq | NumDF | DenDF | F value | Pr(>F) |
| --- | --- | --- | --- | --- | --- | --- |
| *Overyielding* | 0.035 | 0.0025 | 14 | 359.4 | 1.57 | 0.084 |
| *CE* | 0.03 | 0.0021 | 14 | 360 | 1.1 | 0.35 |
| *SE* | 0.0033 | 0.00024 | 14 | 366 | 2.7 | 0.0007 *** |
| *Overperformance in protein content* | 20.64 | 1.47 | 14 | 358.25 | 1.4 | 0.15 |

**Table S7**: Type-III Analysis of Variance Tables of overyielding, Complementarity Effects (CE), Selection Effects (SE), and overperformance in protein content in response to presence/absence of each variety.

*SumSQ,* sum of squares*, MeanSQ,* mean square of error*, DenDF*, degrees of freedom of error term; *NumDF*, degrees of freedom of term; *F-value*, variance ratio; p-values in bold are signiﬁcant at α = 0.05; * (P < 0.05), ** (P < 0.01), *** (P < 0.001). n = 401.

| **Overyielding** | | Sum Sq | | Mean Sq | | NumDF | | DenDF | | F value | | Pr(>F) |
| --- | --- | --- | --- | --- | --- | --- | --- | --- | --- | --- | --- | --- |
| *CH111.16373* | | 0.0014 | | 0.0014 | | 1 | | 368.27 | | 0.863 | | 0.353 |
| *CH211.14074* | | 0.0004 | | 0.0004 | | 1 | | 368.33 | | 0.268 | | 0.605 |
| *Colmetta* | | 0.0102 | | 0.0102 | | 1 | | 368.16 | | 6.397 | | 0.012 * |
| *Campanile* | | 0.0049 | | 0.0049 | | 1 | | 368.39 | | 3.054 | | 0.081 . |
| *Bodeli* | | 0.0000 | | 0.0000 | | 1 | | 368.49 | | 0.002 | | 0.964 |
| *Molinera* | | 0.0004 | | 0.0004 | | 1 | | 368.34 | | 0.262 | | 0.609 |
| *Schilthorn* | |  | |  | |  | |  | |  | |  |
| *Falotta* | |  | |  | |  | |  | |  | |  |
| **CE** | Sum Sq | | Mean Sq | | NumDF | | DenDF | | F value | | Pr(>F) | |
| *CH111.16373* | 0.001 | | 0.001 | | 1 | | 367.99 | | 0.32 | | 0.58 | |
| *CH211.14074* | 0.00001 | | 0.00001 | | 1 | | 368.06 | | 0.005 | | 0.94 | |
| *Colmetta* | 0.01 | | 0.01 | | 1 | | 367.89 | | 4.56 | | 0.03 * | |
| *Campanile* | 0.001 | | 0.001 | | 1 | | 368.11 | | 0.74 | | 0.39 | |
| *Bodeli* | 0.00003 | | 0.00003 | | 1 | | 368.21 | | 0.01 | | 0.91 | |
| *Molinera* | 0.0001 | | 0.0001 | | 1 | | 368.07 | | 0.04 | | 0.85 | |
| *Schilthorn* |  | |  | |  | |  | |  | |  | |
| *Falotta* |  | |  | |  | |  | |  | |  | |
| **SE** | Sum Sq | | Mean Sq | | NumDF | | DenDF | | F value | | Pr(>F) | |
| *CH111.16373* | 3.97E-05 | | 3.97E-05 | | 1 | | 8.09 | | 0.41 | | 0.54 | |
| *CH211.14074* | 1.36E-07 | | 1.36E-07 | | 1 | | 8.13 | | 0.001 | | 0.97 | |
| *Colmetta* | 2.04E-04 | | 2.04E-04 | | 1 | | 8.05 | | 2.11 | | 0.18 | |
| *Campanile* | 3.83E-05 | | 3.83E-05 | | 1 | | 8.23 | | 0.40 | | 0.55 | |
| *Bodeli* | 1.76E-05 | | 1.76E-05 | | 1 | | 8.13 | | 0.18 | | 0.68 | |
| *Molinera* | 2.88E-07 | | 2.88E-07 | | 1 | | 8.09 | | 0.003 | | 0.96 | |
| *Schilthorn* |  | |  | |  | |  | |  | |  | |
| *Falotta* |  | |  | |  | |  | |  | |  | |

| **Overperformance in protein content** | Sum Sq | Mean Sq | NumDF | DenDF | F value | Pr(>F) |
| --- | --- | --- | --- | --- | --- | --- |
| *CH111.16373* | 0.98 | 0.98 | 1 | 366 | 0.93 | 0.336 |
| *CH211.14074* | 0.57 | 0.57 | 1 | 366 | 0.53 | 0.46 |
| *Colmetta* | 0.28 | 0.28 | 1 | 366 | 0.26 | 0.61 |
| *Campanile* | 1.18 | 1.18 | 1 | 366 | 1.11 | 0.29 |
| *Bodeli* | 1.42 | 1.42 | 1 | 366 | 1.34 | 0.24 |
| *Molinera* | 0.74 | 0.74 | 1 | 366 | 0.7 | 0.40 |
| *Schilthorn* |  |  |  |  |  |  |
| *Falotta* |  |  |  |  |  |  |

**Table S8**: Type-III Analysis of Variance Tables of overyielding, Complementarity Effects (CE) and Selection Effects (SE) in response to YES-NO parameter for density plasticity (YES-YES: both varieties increase their relative densities in mixtures compared to their pure stands; YES-NO: one variety increases its density in mixture while the other ones decreases; NO-NO: both varieties decrease their relative densities in mixtures).

*SumSQ,* sum of squares*, MeanSQ,* mean square of error*, DenDF*, degrees of freedom of error term; *NumDF*, degrees of freedom of term; *F-value*, variance ratio; p-values in bold are signiﬁcant at α = 0.05; * (P < 0.05), ** (P < 0.01), *** (P < 0.001). n = 401.

|  | Sum Sq | Mean Sq | NumDF | DenDF | F value | Pr(>F) |  |
| --- | --- | --- | --- | --- | --- | --- | --- |
| **Overyielding** |  |  |  |  |  |  |  |
| *YN density plasticity* | 0.009 | 0.0049 | 2 | 372.1 | 6.03 | 0.0026 | ** |
| **CE** |  |  |  |  |  |  |  |
| *YN density plasticity* | 0.011 | 0.0055 | 2 | 371.4 | 5.93 | 0.0029 | ** |
| **SE** |  |  |  |  |  |  |  |
| *YN density plasticity* | 5.12E-05 | 2.56E-05 | 2 | 373.7 | 0.22 | 0.79 |  |

**Figure S7.** Correlation between Complementarity Effects *sensu* Hector & Loreau (CE, kg/0.45m^2^) and overyielding (kg/0.45m^2^) across all plots (all sites, years, and variety combinations). One data point represents one mixture plot. n = 401.

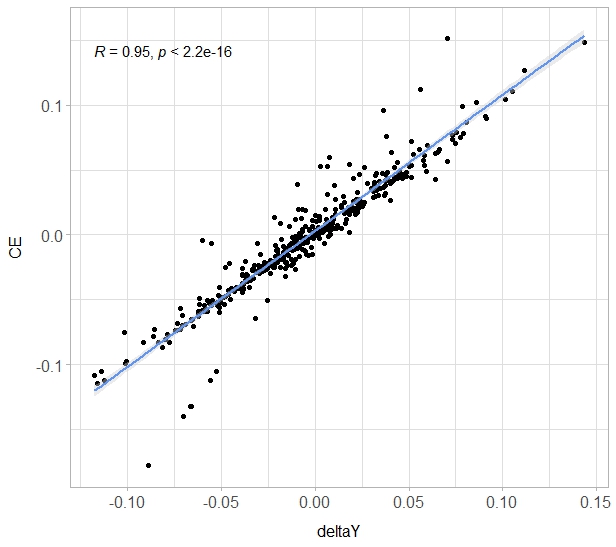

**Table S9**: Regression results from the Structural Equation Modeling with all sites and years (n = 401). Abbreviations: plast = plasticity, b = awned variety, nb = unawned variety, pmg = thousand kernel weight, prot = protein content, deltaY = overyielding, overprot = overperformance in protein content

|  | Estimate | Std.Err | z-value | P(>\|z\|) | Std.lv | Std.all |
| --- | --- | --- | --- | --- | --- | --- |
| **biomass.plast.b ~** |  |  |  |  |  |  |
| dens.plast.b | 0.64 | 0.04 | 17 | < 0.0001 | 0.64 | 0.67 |
| dens.plast.nb | -0.133 | 0.038 | -4 | < 0.0001 | -0.13 | -0.14 |
| **biomass.plast.nb ~** |  |  |  |  |  |  |
| dens.plast.b | -0.1080 | 0.0330 | -3 | < 0.0001 | -0.11 | -0.10 |
| dens.plast.nb | 0.802 | 0.032 | 24.999 | < 0.0001 | 0.802 | 0.794 |
| **pmg.plast.b ~** |  |  |  |  |  |  |
| biomass.plst.b | 10.108 | 3.463 | 2.919 | 0.004 | 10.108 | 0.222 |
| dens.plast.b | -11.096 | 3.348 | -3.315 | 0.001 | -11.096 | -0.253 |
| dens.plast.nb | -7.08E+00 | 2.49E+00 | -2.847 | < 0.0001 | -7.08 | -0.17 |
| **pmg.plast.nb ~** |  |  |  |  |  |  |
| biomss.plst.nb | 8.10E+00 | 3.41E+00 | 2.379 | 0.02 | 8.10 | 0.23 |
| dens.plast.b | -3.71E+00 | 2.13E+00 | -1.741 | 0.08 | -3.71 | -0.10 |
| dens.plast.nb | -7.93E+00 | 3.41E+00 | -2.325 | 0.02 | -7.93 | -0.22 |
| **yield.plast.b ~** |  |  |  |  |  |  |
| dens.plast.b | 0.327 | 0.022 | 14.581 | < 0.0001 | 0.327 | 0.522 |
| biomass.plst.b | 0.201 | 0.023 | 8.779 | < 0.0001 | 0.201 | 0.31 |
| pmg.plast.b | 0.003 | < 0.0001 | 8.541 | < 0.0001 | 0.003 | 0.207 |
| dens.plast.nb | -0.098 | 0.017 | -5.87 | < 0.0001 | -0.098 | -0.16 |
| **yield.plast.nb ~** |  |  |  |  |  |  |
| dens.plast.b | -0.107 | 0.021 | -5.046 | < 0.0001 | -0.107 | -0.154 |
| dens.plast.nb | 0.348 | 0.034 | 10.319 | < 0.0001 | 0.348 | 0.514 |
| biomss.plst.nb | 0.174 | 0.033 | 5.196 | < 0.0001 | 0.174 | 0.259 |
| pmg.plast.nb | 0.004 | 0.001 | 7.856 | < 0.0001 | 0.004 | 0.214 |
| **prot.plast.b ~** |  |  |  |  |  |  |
| biomass.plst.b | -3.065 | 1.792 | -1.711 | 0.087 | -3.065 | -0.122 |
| pmg.plast.b | -0.207 | 0.026 | -7.817 | < 0.0001 | -0.207 | -0.375 |
| dens.plast.b | 1.901 | 1.73 | 1.099 | 0.272 | 1.901 | 0.078 |
| dens.plast.nb | 3.36 | 1.276 | 2.634 | 0.008 | 3.36 | 0.142 |
| **prot.plast.nb ~** |  |  |  |  |  |  |
| biomss.plst.nb | 1.369 | 1.723 | 0.795 | 0.427 | 1.369 | 0.074 |
| pmg.plast.nb | -0.132 | 0.026 | -5.103 | < 0.0001 | -0.132 | -0.252 |
| dens.plast.nb | -0.169 | 1.728 | -0.098 | 0.922 | -0.169 | -0.009 |
| dens.plast.b | 4.011 | 1.076 | 3.73 | < 0.0001 | 4.011 | 0.21 |
| **deltaY ~** |  |  |  |  |  |  |
| yield.plast.b | 1.026 | 0.007 | 151.702 | < 0.0001 | 1.026 | 1.067 |
| yield.plast.nb | 0.999 | 0.006 | 169.782 | < 0.0001 | 0.999 | 1.15 |
| biomass.plst.b | -0.003 | 0.004 | -0.76 | 0.447 | -0.003 | -0.005 |
| biomss.plst.nb | -0.001 | 0.004 | -0.261 | 0.794 | -0.001 | -0.002 |
| prot.plast.b | < 0.0001 | < 0.0001 | -0.417 | 0.677 | < 0.0001 | -0.002 |
| prot.plast.nb | < 0.0001 | < 0.0001 | -0.202 | 0.84 | < 0.0001 | -0.001 |
| **overprot.ini ~** |  |  |  |  |  |  |
| yield.plast.b | 6.803 | 1.921 | 3.541 | < 0.0001 | 6.803 | 0.273 |
| yield.plast.nb | 0.11 | 1.672 | 0.066 | 0.948 | 0.11 | 0.005 |
| biomass.plst.b | -1.442 | 1.12 | -1.287 | 0.198 | -1.442 | -0.089 |
| biomss.plst.nb | 1.118 | 1.075 | 1.04 | 0.298 | 1.118 | 0.074 |
| prot.plast.b | 0.299 | 0.03 | 10.007 | < 0.0001 | 0.299 | 0.465 |
| prot.plast.nb | 0.25 | 0.038 | 6.545 | < 0.0001 | 0.25 | 0.305 |

Table S9: Regression results from the Structural Equation Modeling with all only Changins (n = 135).

Abbreviations: plast = plasticity, b = awned variety, nb = unawned variety, pmg = thousand kernel weight, prot = protein content, deltaY = overyielding, overprot = overperformance in protein content

|  | Estimate | Std.Err | z-value | P(>\|z\|) | Std.lv | Std.all |
| --- | --- | --- | --- | --- | --- | --- |
| **biomass.plast.b ~** |  |  |  |  |  |  |
| dens.plast.b | 0.543 | 0.091 | 5.963 | 0 | 0.543 | 0.493 |
| dens.plast.nb | -0.101 | 0.076 | -1.322 | 0.186 | -0.101 | -0.109 |
| **biomass.plast.nb** ~ |  |  |  |  |  |  |
| dens.plast.b | -0.149 | 0.049 | -3.063 | 0.002 | -0.149 | -0.134 |
| dens.plast.nb | 0.806 | 0.041 | 19.789 | 0 | 0.806 | 0.864 |
| **height.plast.b ~** |  |  |  |  |  |  |
| biomass.plst.b | 3.462 | 8.438 | 0.41 | 0.682 | 3.462 | 0.044 |
| biomss.plst.nb | 29.747 | 15.608 | 1.906 | 0.057 | 29.747 | 0.379 |
| dens.plast.b | 18.03 | 9.362 | 1.926 | 0.054 | 18.03 | 0.207 |
| dens.plast.nb | -10.409 | 14.16 | -0.735 | 0.462 | -10.409 | -0.142 |
| **height.plast.nb ~** |  |  |  |  |  |  |
| biomss.plst.nb | 61.577 | 14.572 | 4.226 | 0 | 61.577 | 0.808 |
| dens.plast.b | 17.29 | 8.826 | 1.959 | 0.05 | 17.29 | 0.204 |
| dens.plast.nb | -37.325 | 13.311 | -2.804 | 0.005 | -37.325 | -0.525 |
| biomass.plst.b | -1.771 | 7.856 | -0.225 | 0.822 | -1.771 | -0.023 |
| **pmg.plast.b ~** |  |  |  |  |  |  |
| biomass.plst.b | 10.27 | 5.067 | 2.027 | 0.043 | 10.27 | 0.209 |
| dens.plast.b | -10.69 | 5.769 | -1.853 | 0.064 | -10.69 | -0.198 |
| dens.plast.nb | -5.576 | 4.311 | -1.293 | 0.196 | -5.576 | -0.123 |
| height.plast.b | -0.065 | 0.056 | -1.16 | 0.246 | -0.065 | -0.106 |
| **pmg.plast.nb ~** |  |  |  |  |  |  |
| biomss.plst.nb | -3.589 | 7.079 | -0.507 | 0.612 | -3.589 | -0.11 |
| dens.plast.b | -2.3 | 3.67 | -0.627 | 0.531 | -2.3 | -0.063 |
| dens.plast.nb | -0.806 | 6.237 | -0.129 | 0.897 | -0.806 | -0.026 |
| height.plst.nb | -0.023 | 0.044 | -0.519 | 0.604 | -0.023 | -0.054 |
| **yield.plast.b ~** |  |  |  |  |  |  |
| dens.plast.b | 0.333 | 0.042 | 7.991 | 0 | 0.333 | 0.555 |
| biomass.plst.b | 0.096 | 0.038 | 2.55 | 0.011 | 0.096 | 0.176 |
| pmg.plast.b | 0.001 | 0.001 | 1.963 | 0.05 | 0.001 | 0.119 |
| dens.plast.nb | -0.159 | 0.031 | -5.19 | 0 | -0.159 | -0.316 |
| height.plast.b | -0.001 | 0 | -1.773 | 0.076 | -0.001 | -0.107 |
| **yield.plast.nb ~** |  |  |  |  |  |  |
| dens.plast.b | -0.097 | 0.037 | -2.629 | 0.009 | -0.097 | -0.127 |
| dens.plast.nb | 0.263 | 0.063 | 4.166 | 0 | 0.263 | 0.411 |
| biomss.plst.nb | 0.323 | 0.072 | 4.499 | 0 | 0.323 | 0.47 |
| pmg.plast.nb | 0.002 | 0.001 | 2.207 | 0.027 | 0.002 | 0.094 |
| height.plst.nb | 0 | 0 | -0.124 | 0.901 | 0 | -0.006 |
| **prot.plast.b ~** |  |  |  |  |  |  |
| biomass.plst.b | 1.225 | 2.522 | 0.486 | 0.627 | 1.225 | 0.05 |
| pmg.plast.b | -0.106 | 0.041 | -2.613 | 0.009 | -0.106 | -0.212 |
| dens.plast.b | -0.55 | 2.777 | -0.198 | 0.843 | -0.55 | -0.02 |
| height.plast.b | 0.069 | 0.027 | 2.571 | 0.01 | 0.069 | 0.221 |
| dens.plast.nb | 4.121 | 2.043 | 2.017 | 0.044 | 4.121 | 0.181 |
| **prot.plast.nb ~** |  |  |  |  |  |  |
| biomss.plst.nb | 0.397 | 4.72 | 0.084 | 0.933 | 0.397 | 0.017 |
| pmg.plast.nb | -0.062 | 0.055 | -1.125 | 0.261 | -0.062 | -0.089 |
| dens.plast.nb | -2.305 | 4.15 | -0.555 | 0.579 | -2.305 | -0.109 |
| height.plst.nb | 0.078 | 0.027 | 2.908 | 0.004 | 0.078 | 0.263 |
| dens.plast.b | 6.093 | 2.381 | 2.559 | 0.01 | 6.093 | 0.242 |
| **deltaY ~** |  |  |  |  |  |  |
| yield.plast.b | 1.007 | 0.011 | 92.41 | 0 | 1.007 | 0.886 |
| yield.plast.nb | 0.983 | 0.013 | 73.894 | 0 | 0.983 | 1.1 |
| biomass.plst.b | 0.001 | 0.005 | 0.111 | 0.911 | 0.001 | 0.001 |
| biomss.plst.nb | 0.005 | 0.009 | 0.573 | 0.567 | 0.005 | 0.009 |
| prot.plast.b | 0 | 0 | -1.704 | 0.088 | 0 | -0.014 |
| prot.plast.nb | 0 | 0 | -1.268 | 0.205 | 0 | -0.011 |
| **overprot.ini ~** |  |  |  |  |  |  |
| yield.plast.b | 15.045 | 3.37 | 4.465 | 0 | 15.045 | 0.464 |
| yield.plast.nb | -6.875 | 4.174 | -1.647 | 0.1 | -6.875 | -0.269 |
| biomass.plst.b | -3.351 | 1.668 | -2.009 | 0.045 | -3.351 | -0.189 |
| biomss.plst.nb | 5.342 | 2.907 | 1.838 | 0.066 | 5.342 | 0.305 |
| prot.plast.b | 0.239 | 0.063 | 3.828 | 0 | 0.239 | 0.333 |
| prot.plast.nb | 0.111 | 0.072 | 1.549 | 0.121 | 0.111 | 0.143 |

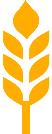

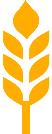

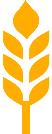

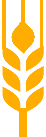

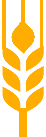

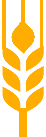

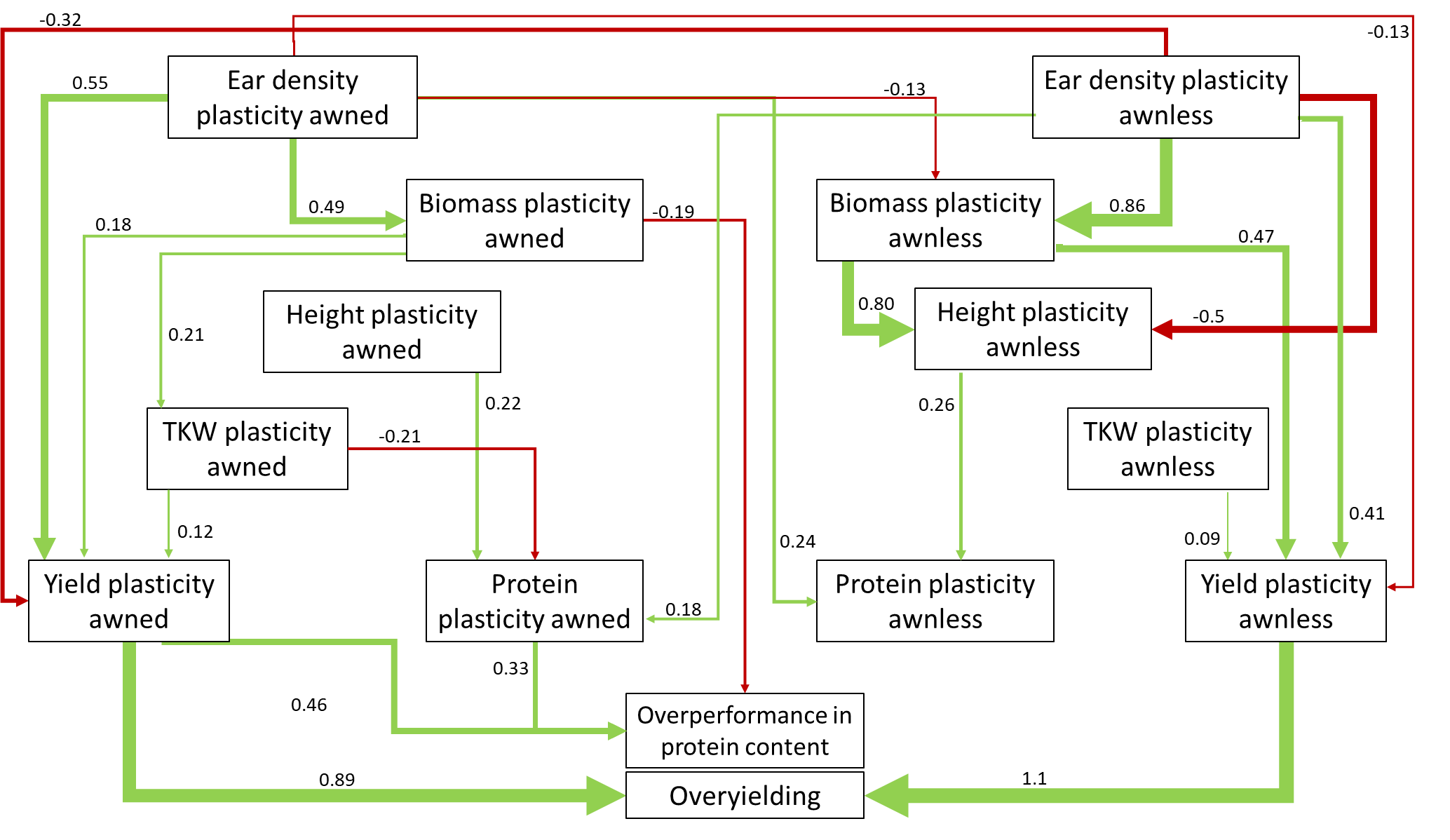

**Figure S9:** Ear density (a) and grain yield (b) of the varieties in pure stands. Dots represent the mean values across replicates, sites, and years; lines represent the standard error.

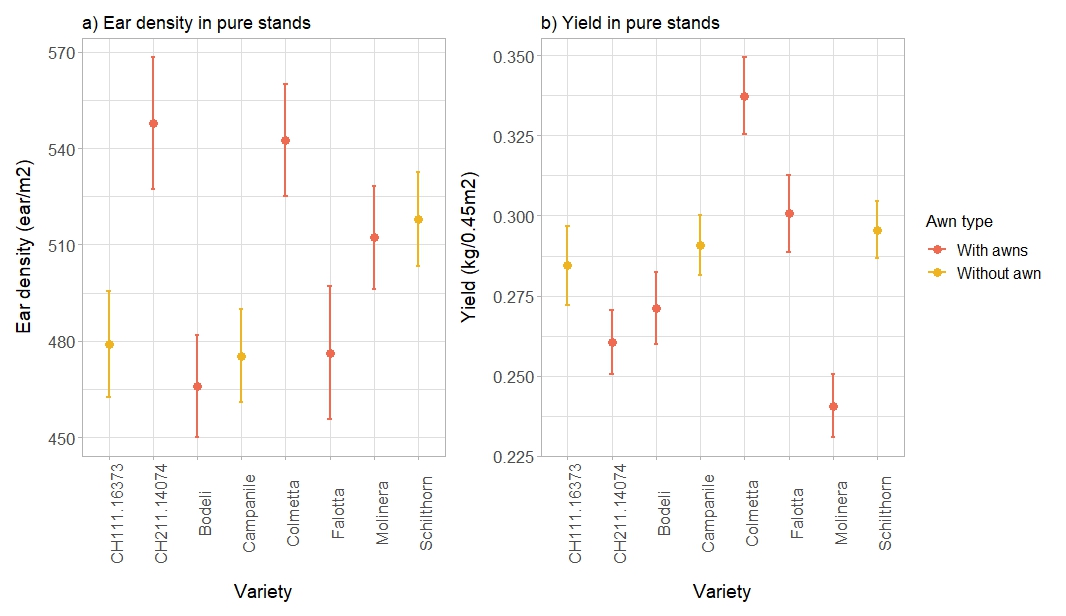

**Figure S10:** Adjusted coefficients of variation of density (a) and grain yield (b) in pure stands. Adjusted coefficients of variation were calculated per variety across years and sites.

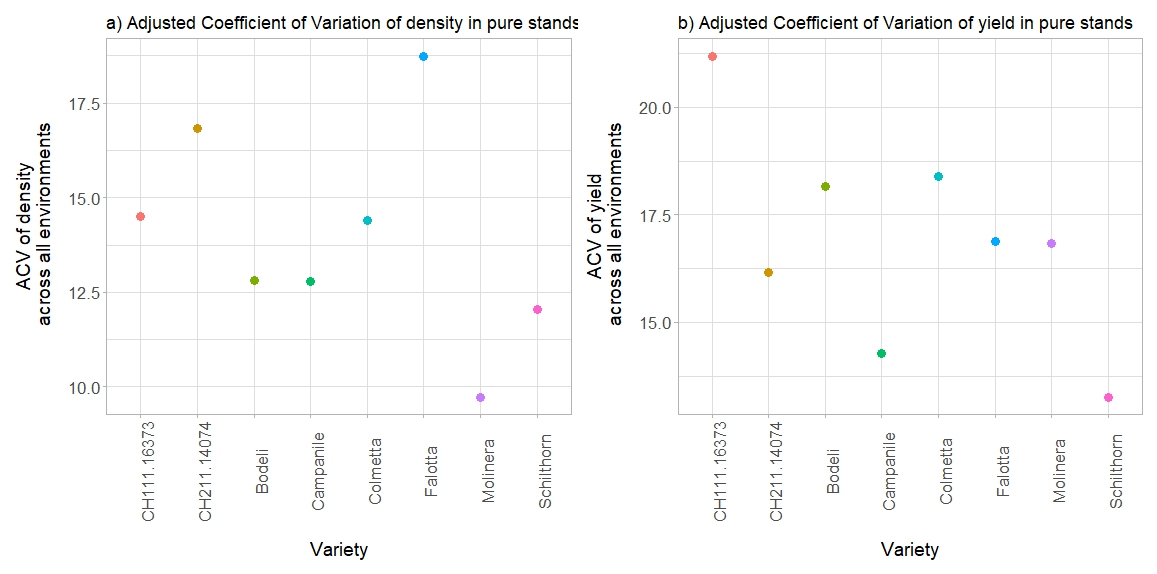

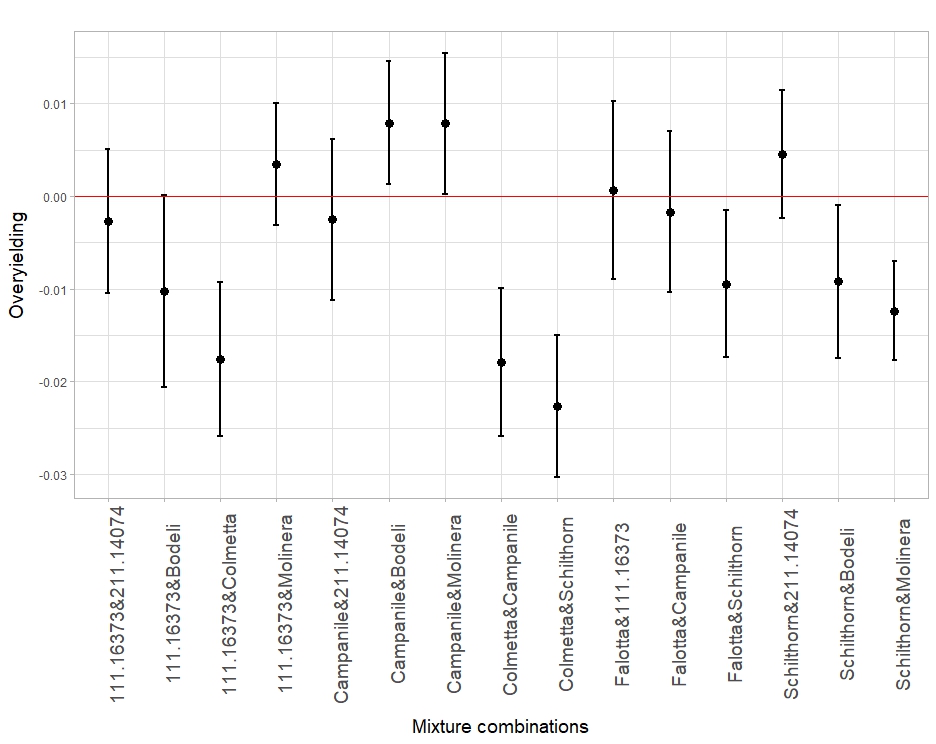

**Figure S11:** Overyielding (kg/0.45m^2^) per mixture combination, averaged across years and sites. Dots represent the mean values across plots; lines represent the standard error.
